# Epigenetic plasticity via adaptive DNA hypermethylation and clonal expansion underlie resistance to oncogenic pathway inhibition in pancreatic cancer

**DOI:** 10.1101/2022.05.20.492826

**Authors:** Laura K. Godfrey, Jan Forster, Sven-Thorsten Liffers, Christopher Schröder, Johannes Köster, Leonie Henschel, Kerstin U. Ludwig, Marija Trajkovic-Arsic, Diana Behrens, Aldo Scarpa, Rita T. Lawlor, Kathrin E. Witzke, Barbara Sitek, Steven A. Johnsen, Sven Rahmann, Bernhard Horsthemke, Michael Zeschnigk, Jens T. Siveke

## Abstract

Pancreatic ductal adenocarcinoma (PDAC) is an aggressive cancer with poor prognosis. Drug resistance is the major cause for therapeutic failure in PDAC patients with progressive disease. The mechanisms underlying resistance formation are complex and remain poorly understood.

To gain insights into molecular changes during the formation of resistance to oncogenic MAPK pathway inhibition we utilized short-term passaged primary tumor cells from ten PDACs of genetically engineered mice. We followed gain and loss of resistance upon MEK_i_ exposure and withdrawal by longitudinal integrative analysis of whole genome sequencing, whole genome bisulfite sequencing, RNA-sequencing and mass spectrometry data.

We found that resistant cell populations under increasing MEK_i_ treatment evolved by the expansion of a single clone but were not a direct consequence of known resistance-conferring mutations. Rather, resistant cells showed adaptive DNA hypermethylation of 209 and hypomethylation of 8 genomic sites, most of which overlap with regulatory elements known to be active in murine PDAC cells. Both DNA methylation changes and MEK_i_ resistance were transient and reversible upon drug withdrawal. The effector caspase *CASP3* is one of the 114 genes for which transcriptional downregulation inversely correlated with the methylation status of the associated DNA region. CASP3 inactivation in resistant cells led to attenuation of drug-induced apoptosis which could be reversed by DNA methyltransferase inhibition with remarkable sensitivity exclusively in the resistant cells.

Overall, our data provide a context for characterization and targeting of epigenetically mediated resistance mechanisms in PDAC.

## Introduction

Overcoming treatment resistance is a critical challenge for improving the prognosis for patients with pancreatic ductal adenocarcinoma (PDAC). Genetically, only four non-targetable genes (*KRAS*, *TP53*, *CDKN2A*, *SMAD4*) are known to be recurrently mutated in PDAC so far. Many low frequency alterations found in various genes reflect a remarkable inter- and intra-individual tumor heterogeneity(1–3). Despite major advances in defining the genomic landscape of PDAC, the inherent and acquired resistance mechanisms during tumor evolution and upon therapeutic perturbations remain a considerable challenge.

Oncogenic *KRAS* mutations represent a therapeutic target in PDAC. While direct inhibitors for the most frequent KRAS G12D and G12V variants are still lacking, potent and specific inhibitors for the downstream effector mitogen-activated protein kinase (MAPK) signaling pathway exist, including highly selective inhibitors against MEK, a component of the MAPK pathway(4). However, despite promising results in preclinical model systems(5–7), MEK inhibitors (MEK_i_) have failed in clinical trials due to rapid induction of resistance(8–11). Several cellular processes including mitochondrial function, nucleotide synthesis, protective autophagy or the deregulation of YAP, SHP or ERBB have been reported to be involved in MEKi resistance(12–19). However, their clinical relevance and the underlying regulatory circuits still remain to be identified.

Growing evidence supports a concept where tumor cells utilize epigenetic mechanisms to adapt to varying conditions, including MEK_i_ treatment of PDAC(20–22). Mutations affecting epigenetic readers and writers such as enzymes controlling histone modifications and DNA methylation are frequently found in PDAC and other cancers(2,23,24).

In this study, we focus on longitudinal characterization of molecular alterations underlying MEK_i_ resistance. We combine multi-omics technologies on the genetic, epigenetic, transcriptomic and protein levels in primary genetically engineered mouse model (GEMM)-derived PDAC cells, thereby minimizing interindividual genetic and epigenetic heterogeneity typically confounding patient-derived tumor analyses. We address adaptive epigenetic changes under therapeutic pressure and identify a vulnerability of MEK_i_-induced resistant PDAC cells to DNA methyltransferase inhibitors (DNMT_i_). We found adaptive DNA hypermethylation in cells that acquired MEK_i_ resistance and characterized its dynamics upon drug withdrawal.

## Methods

### Generation of primary murine PDAC cell lines

Tumor pieces derived from *Ptf1a^wt/Cre^; Kras^wt/LSL-G12D^; Trp53^loxP/loxP^* mice were incubated at 37°C and 5% CO_2_ in high-glucose Dulbecco’s Modified Eagle’s Medium (DMEM) (Thermo Fisher Scientific) containing 10% fetal bovine serum (Thermo Fisher Scientific), 1% penicillin/streptomycin (Thermo Fisher Scientific), and 1% non-essential amino acids (Sigma-Aldrich) until tumor cells emigrated. PCR-based mycoplasma testing was performed on a regular basis. Cell lines #1, #3, #4, #5 and #8 were derived from male mice and cell lines #2, #6, #7, #9 and #10 from female mice. All cell lines used are available from the corresponding author upon reasonable request.

### MEK_i_ resistance induction in primary murine PDAC cell line

MEK_i_ resistance was induced in ten different low-passage cell lines (< 4-12 passages). Therefore, cells were treated with increasing doses of trametinib (LKT) until they grew in 100x of their IC50 (800 nM to 4200 nM trametinib). One batch of each cell line was cultivated with 100x IC50 of trametinib in the culture medium (termed resistant hereafter) with medium exchange every 2-3 days on a regular basis. Another batch was kept under drug withdrawal and samples were named according to their passage number in drug-free medium (Px).

### Cell viability assays

Cell viability assays were performed with four to six different cell lines. Cell numbers were optimized for 80% confluency in 96- or 384-well plates, respectively. Drugs targeting different epigenetic modifiers (trametinib (LKT), decitabine (Sigma-Aldrich), JQ1 (Cayman), suberoylanilide hydroxamic acid (SAHA) (Selleckchem), mocetinostat (Selleckchem) dissolved in dimethyl sulfoxide (DMSO) (Sigma-Aldrich) were printed in the indicated logarithmic concentration ranges using the D300e Digital Dispenser (Tecan). The DMSO concentration in each well was adjusted to the highest value on the plate which was set to < 0.1% of the assay volume. Sealed plates were frozen at -80°C until use.

Cells were detached by 0.05% trypsin - ethylenediamine tetraacetic acid (EDTA) (1x) (Thermo Fisher Scientific) and recovered by centrifugation. Optimized cell numbers for 80% confluency at the end of experiment were seeded with the Multidrop Combi Dispenser (Thermo Fisher Scientific) onto the pre-printed plates and incubated at 37°C and 5% CO_2_.

Cell viability was determined using the CellTiter-Glo® Luminescent Cell Viability Assay (Promega) according to manufacturer’s instruction. The luminescence signal was measured with a Tecan Spark® 10 M multiplate reader (Tecan) for 500 ms.

Data were normalized to the signal of DMSO treated cells. IC50 determination was performed using the Graph Pad Prims v. 7.03 ‘log (inhibitor) vs. response (three parameters)’ equation.

Synergism was analyzed by applying the Loewe (25) method implemented in Combenefit v. 2.02 (26).

### Extraction of total protein and Simple Western analysis

For total protein isolation, RIPA buffer (Cell Signaling Technology) containing protease- and phosphatase-inhibitor cocktails (Sigma-Aldrich) was used to lyse Dulbecco’s phosphate-buffered saline (DPBS)-washed (Thermo Fisher Scientific) cell pellets on ice for 20 min. To remove debris, lysates were centrifuged at 4°C and full-speed for 10 min. Afterwards, the protein concentration was determined with the Pierce BCA Protein Assay Kit (Thermo Fisher Scientific).

If nothing else indicated 0.2 mg/ml protein per 12-230 kDa capillary were used in a Simple Western analysis using the Wes instrument (ProteinSimple) as suggested by the manufacturer’s protocol. Antibodies against the following proteins were used in the indicated dilutions: ERK1/2 (#4695, RRID:AB_390779, Cell Signaling Technology, 1:50), JUN (#9165, RRID:AB_2130165, Cell Signaling Technology, 1:50), p-JUN (#9164, RRID:AB_330892, Cell Signaling Technology, 1:5), p-ERK1/2 (#4376, RRID:AB_331772, Cell Signaling Technology, 1:15), Vinculin (#13901, RRID:AB_2728768, Cell Signaling Technology, 1:30,000).

### CASP3 activity assay

CASP3 activity was assessed in 96-well plates using duplicates for control conditions and triplicates for drug treatments. Parental and resistant cells of three different lines were tested. Compounds were pre-printed with the D300e Digital Dispenser (Tecan), normalized for DMSO content. The following drug concentrations were used: trametinib (LKT) 100x IC50 of each individual cell line, decitabine (Sigma-Aldrich) 0.5 µM, trametinib 0.3 µM + decitabine 0.5 µM. Optimized cell numbers for each cell line to reach 80% confluency at the end of experiment were seeded and incubated for the indicated time points. The Caspase-Glo® Assay (Promega) was applied according to manufacturer’s instructions. It contains a CASP3/7 specific luminogenic substrate and luminescence was detected with a Tecan Spark® 10 M multiplate reader (Tecan). Signals were corrected for viable cells measured by CellTiter-Glo® Luminescent Cell Viability Assay (Promega) as described in ‘Cell viability assays’.

### Flow cytometric quantification of cell death

Appropriate cell numbers to reach 80% confluency at the end of experiment were individually determined for each of the three tested cell lines and seeded in 12-well plates pre-printed with the following drug concentrations: trametinib (LKT) 100x IC50 in each individual cell line, decitabine (Sigma-Aldrich) 0.5 µM, trametinib 0.3 µM + decitabine 0.5 µM (D300e Digital Dispenser (Tecan). DMSO concentrations per well were normalized to the highest concentration on the plate. After incubation for 84 h, cells were detached by accutase (Sigma-Aldrich) and stained with the FITC Annexin V Apoptosis Detection Kit I (BD Bioscience) using Annexin V-FITC and PI both 1:40 for 15 min at room temperature protected from light. Fluorescence was analyzed by flow cytometry on a FACS Aria system (BD Bioscience). The percentage of each cell population was determined with the FlowJo software v. 10.5.3 (Becton, Dickinson and Company).

### Cytogenetic analysis

Parental cells of cell lines #3 and #9 were treated with colcemid for 4 h prior to harvest. Culture solution was centrifuged, the cell pellet was resuspended in a hypotonic 75 mM KCl-solution and incubated for 20 min at 37°C. After centrifugation, cells were resuspended by dropwise adding 8 ml of an ice-cold fixative solution (3:1 mixture methanol and acetic acid). Cells were washed 3 times in 8 ml ice-cold fixative solution for 10 min each and dropped onto a fat-free and watered glass slide that was then air-dried overnight at 60°C. Chromosomes were stained with Giemsa and examined under the microscope using 140x magnification.

### Patient-derived xenografts

All mice experiments were carried out by D. Behrens at EPO GmbH, Berlin-Buch, and performed according to the German Animal Protection Law with approval from the responsible authorities. The *in vivo* procedures were consistent and in compliance with the UKCCCR guidelines. Already established patient-derived xenografts of pancreatic adenocarcinoma from three different male patients at passage number 2 were received from ARC-NET, University of Verona. The materials used have been collected under Program 1885 protocol 52438 on 23/11/2010 and Program 2172 protocol 26773CE 23/05/2012. The protocols include informed consent of the patients and were approved by the local ethics committee of the Integrated University Hospital Trust of Verona. At the time of surgery patient 1 was 59 years old, patient 2 65 years and patient 3 53 years. Mice were maintained in the pathogen-free animal facility following institutional guidelines and with approval from the responsible authorities. The animals were housed under pathogen-free conditions in individually ventilated cages under standardized environmental conditions (22°C room temperature, 50 ± 10% relative humidity, 12 hours light-dark rhythm). They received autoclaved food and bedding (Ssniff) and acidified (pH 4.0) drinking water *ad libitum*.

Tumor pieces of 3 mm^3^ were transplanted subcutaneously into NOD/SCID-mice with knocked IL2γ receptor (NSG mice) within 24 h after explant from donor mice. Remaining tumor tissue was preserved in DMSO or snap-frozen for later propagation or analyses. Engrafted tumors at a size of about 1 cm^3^ were surgically excised and fragments of 2 - 3 mm^3^ re-transplanted into immune deficient NMRI:nu/nu mice for further passage. Tumors were passaged not more than 10 times.

For drug screening studies tumor material was implanted subcutaneously into appropriate cohorts of NMRI:nu/nu mice (n=3 per treatment group). At advanced tumor size (200 mm³), mice were randomized and treated with 1 mg/kg trametinib (p.o., daily), 0.2 mg/kg decitabine (s.c., three times weekly). To further mouse cohorts the combinations of trametinib and decitabine were applied. Tumor size was measured with a caliper instrument and monitored during the entire experiment with the measurements of two perpendicular tumor diameters using the spheroid equation: tumor volume = [(tumor width)^2^ x tumor length] x 0.5. Treatment was continued over a period of two weeks unless tumor size exceeded 10% of animal body weight or animals showed loss of more than 15% body weight. Six hours after last treatment animals were sacrificed and tumor samples preserved for further analyses.

### Mass spectrometry

#### Sample preparation

Cell pellets of parental, resistant and P12 cells of six different lines were resuspended in 100 μl 50 mM ammonium bicarbonate and 0.1% sodium deoxycholate (NaDOC) for cell lysis. Samples were sonicated on ice for 10 min and centrifuged (16,000 g, 15 min, 4°C). Protein concentration was determined via Bradford assay. Due to a very low concentration, technical replicates were pooled. The samples were ridded of remaining viscosity with 10 impulses at 5% power by ultrasonic homogenization via Sonopuls HD 200 MS 72 (Badelin) and centrifuged (16,000 g, 15 min, 4°C). Protein amount was determined via amino acid analysis. DTT (5 mM) was added to the sample for reduction (30 min, 60°C), followed by iodoacetamide (15 mM) for alkylation (30 min, room temperature, in the dark). Lysed proteins were tryptically digested over night at 37°C (trypsin/protein ratio 1/24). For acidification, trifluoroacetic acid (TFA) (0.5%) was added (30 min, 37°C), samples were centrifuged (10 min, 16,000 g) for removal of NaDOC and supernatant transferred to glass vials, dried in a vacuum centrifuge, and dissolved in 0.1% TFA. A sample amount corresponding to 275 ng was used for one liquid chromatography tandem-mass spectrometry (LC-MS/MS) measurement.

#### LC-MS/MS parameters

LC–MS/MS analysis was performed on a LTQ Orbitrap Elite instrument (Thermo Fisher Scientific) coupled online to an upstream-connected Ultimate 3000 RSLCnano high-performance liquid chromatography system (Dionex). Samples were measured in a shuffled manner. Peptides dissolved in 0.1% TFA were pre-concentrated on a C18 trap column (Acclaim PepMap 100; 100 μm × 2 cm, 5 μm, 100 Å; Thermo Fisher Scientific) within 7 min at a flow rate of 30 μl/min with 0.1% TFA. Peptides were then transferred to an in-house packed C18 analytical column (ReproSil®-Pur from Dr. Maisch HPLC GmbH, Ammerbuch, Germany, 75 μm × 40 cm, 1.9 μm, 120 Å). Peptides were separated with a gradient from 5%–40% solvent B over 98 min at 300 nl/min and 65°C (solvent A: 0.1% formic acid; solvent B: 0.1% formic acid, 84% acetonitrile). Full-scan mass spectra in the Orbitrap analyzer were acquired in profile mode at a resolution of 60,000 at 400 m/z and within a mass range of 350 - 2000 m/z. MS/MS spectra were acquired in data-dependent mode at a resolution of 5,400. For MS/MS measurements, the 20 most abundant peptide ions were fragmented by collision-induced dissociation (normalized collision energy (NCE) of 35) and measured for tandem mass spectra in the linear ion trap.

#### Protein identification and quantification

Proteins were identified with Proteome Discoverer v. 1.4 (Thermo Fisher Scientific). Spectra were searched against the UniProtKB/Swiss-Prot database (Release 2018_11; 53,780 entries) using Mascot v. 2.5 (Matrix Science, London, UK). Taxonomy setting was *Mus musculus* and mass tolerance was 5 ppm and 0.4 Da for precursor and fragment ions, respectively. Dynamic and static modifications were considered for methionine (oxidation) and cysteine (carbamidomethyl), respectively. The FDR was calculated with the Proteome Discoverer Target Decoy PSM Validator function, and identifications with a FDR > 1% were rejected. The software Progenesis QI v. 2.0.5387.52102 (Nonlinear Dynamics) was used for label-free quantification. The obtained raw files were aligned to a reference run and a master map of common features was applied to all experimental runs to adjust for differences in retention time. Ion charge states of 2+, 3+, and 4+ with a minimum of three isotope peaks were considered, and raw ion abundances were normalized for automatic correction of technical or experimental variations between runs. Quantified features were identified using the obtained Proteome Discoverer identifications. All non-conflicting peptides were considered for protein quantification.

#### Statistics

Progenesis calculates statistical significance of measured differences (ANOVA p-value) and ratios of means (fold changes). However, when more than two different groups are processed with the software, the resulting p-values only state that a significant difference between two of those groups exists. Similarly, only maximal fold changes are calculated. Application of a post-hoc test was therefore necessary. Normalized protein abundances were obtained from Progenesis and analyzed by applying ANOVA followed by Tukey’s honest significant difference (HSD) method. Fold changes between groups were determined based on normalized abundances while ANOVA was calculated using arcsinh-transformed data for consistency with the Progenesis QI software. The FDR was controlled by adjusting ANOVA p-values using the method of Benjamini and Hochberg (27). For proteins with adjusted ANOVA p-values below the significance level of α=0.05, the TukeyHSD method was applied to further characterize the identified differences in abundance levels between groups. Proteins were considered differentially abundant between groups with a log_2_ fold change ≥ 1 or ≤ -1 and an adjusted p-value < 0.05.

### Isolation of nucleic acids

DNA and RNA were isolated using the Maxwell® RSC Cultured Cells DNA and the Maxwell® RSC simplyRNA Cells Kit (Promega) according to manufacturer’s instruction. Nuclease-free water was used for DNA elution.

### RNA-sequencing

#### Sequencing

RNA-sequencing of parental, resistant and P12 cells of six different cell lines was performed by CeGaT (Tübingen). In addition, P5 cells of cell lines #3, #7 and #9 were sequenced. For library preparation the TruSeq Stranded mRNA Kit (Illumina) was used with 100 ng input RNA and 2x 100 bp were sequenced on a HiSeq 4000 (Illumina) or a NovaSeq 6000 system (Illumina).

#### Read processing and quantification

Demultiplexing of the sequencing reads was performed with Illumina CASAVA v. 2.17 or bcl2fastq v. 2.19. Adapters were trimmed with Skewer v. 0.1.116 or 0.2.2(28).

Transcripts were quantified using the quasi-mapping approach of salmon v. 0.12 (29). TXImport v. 1.6(30) and DESeq2 v. 1.18(31) were used to import transcript-level counts, convert them to gene-level counts and perform differential expression analysis between all four cell states (parental, resistant, P5, P12). Results were multiple test-corrected by the Benjamini-Hochberg method.

#### Principal component analysis

Principal component analysis (PCA) was performed on the normalized gene-level counts of all expressed genes.

#### Hierarchical clustering

Hierarchical clustering of significantly differentially expressed genes (Benjamini-Hochberg adjusted p-value < 0.01; log_2_ fold change > 1 or log_2_ fold change < -1) between parental versus the union of resistant, P5, P12 and resistant versus the union of parental, P5, P12 was computed by the ward.D2 method. Additionally, the same method was used to cluster samples based on PDAssigner genes(32) or PDAC subtype associated genes defined by Bailey et. al.(33) and Moffitt et al.(34).

#### Gene set enrichment analysis

GSEA(35) was performed using default settings and gene set permutation.

#### Score to define reverting Transcripts

In order to identify differentially expressed genes that show a similar expression pattern in parental and P12 samples, a score was defined based on the log_2_ fold change between parental/P12 and resistant samples.

The score was defined as follows:

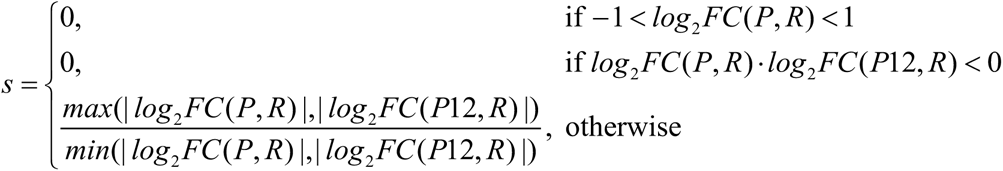

where *log*_2_*FC*(*A,B*) describes the log_2_ fold change between A and B. Genes were only considered if the log_2_ fold change between parental and resistant cell state was reasonably large.

### Whole genome bisulfite sequencing

#### Sequencing and alignment

Whole genome bisulfite sequencing (WGBS) of parental, resistant and P12 cells of cell lines #3 and #9 was performed at the Genomics and Proteomics Core Facility of the German Cancer Research Center (GPCF DKFZ, Heidelberg) using the TruSeq DNA PCR-free Methyl protocol (Illumina) for library preparation. A HiSeq X machine (Illumina) was used for 150 bp paired-end sequencing. Reads were mapped using bwa-meth v. 0.2.0(36) on the GRCm38 assembly with added PhiX genome as a sequencing control.

#### Calculation of methylation levels

CpG methylation levels were computed using an in-house script filtering reads with a mapping quality < 30 and bases with base quality < 17.

#### Differentially methylated region detection

The BSmooth algorithm of bsseq v. 1.10(37) was used to detect DMRs between parental and resistant samples, with every DMR containing a minimum of four CpGs and a minimum difference in methylation level of 0.4.

#### Nearest genes

For each DMR, the nearest flanking genes were determined by finding the nearest TSSs to each region using BEDTools closest v. 2.27(38).

#### Integration with RNA-seq data

Expression changes of the two nearest genes of every DMR were assessed from RNA-sequencing data as described above. Genes with a log_2_ fold change > 1 were defined as upregulated in resistant cells, those with a log_2_ fold change < -1 as downregulated in resistant cells.

#### Genomic regions

The localization of DMRs relative to genes and CpG islands was performed using BEDTools intersect v. 2.27 (Quinlan and Hall, 2010). Reference data were taken from Ensembl build 93(39) (genes) and the UCSC database (CpG islands).

Reference data for shore and shelve regions were created using BEDTools flank v. 2.27(38), shores were defined as regions up to 2000 bp away from CpG islands and shelves as regions up to 2000 bp away from shores.

#### Methylation score to define reverting DMRs

A score was used to model the methylation changes between parental, resistant and P12 cells, where scores > 0.5 indicate that the P12 methylation is closer to the parental level, scores < 0.5 indicate the P12 methylation is closer to the resistant level.

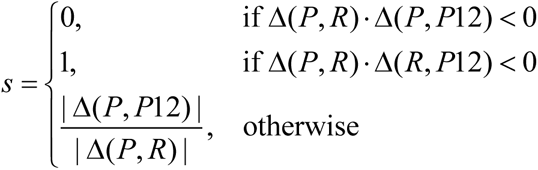

where Δ(*A, B*) describes the difference in methylation between A and B. A cut-off of 0.44 (90% quantile) was used to define DMRs as reverting.

#### Conservation

The UCSC liftOver tool was used together with the mm10ToHg38 liftOver chain in order to identify regions in the human genome that are associated with the murine DMRs. The minimum ratio of bases that need to remap to define a region as valid liftOver was set to 0.5.

#### Regulatory Regions

Overlaps between DMRs and transcription factor binding sites, miRNA target regions and VISTA enhancers were computed using BEDTools intersect v. 2.27(38). Reference data were taken from the Ensembl regulation build 93(39).

The findMotifsGenome script from homer v. 4.9 was used in order to find enrichment of known binding motifs from the homer library(40). The script was used with standard parameters (-size 200 -cpg) on the GRCm38 assembly, comparing reverting DMRs to random background sequences.

#### Overlap with chromatin marks from PDAC organoids

Organoid data available from Roe et al.(41) were used to check whether DMRs overlap with open chromatin or enhancer regions.

Analysis was performed on the following organoids

H3K27ac-ChIP-seq:

- N5, N6 – normal pancreatic organoids(42)
- T3, T6, T19, T23, T33, T34 – tumor organoids

ATAC-seq:

- N5, N6 – normal pancreatic organoids(42)
- T3, T6, T23 – tumor organoids

Reads were aligned with BWA-MEM v. 0.7.17(43) against the GRCm38 genome. Duplicate reads were further removed using Samtools v. 1.9.

Identification of ChIP-seq and ATAC-seq peaks was performed using MACS2 v. 2.1.2(44, 45) callpeak function with default settings. Resulting narrowPeak files were further merged with BEDtools merge v. 2.27(38) and overlapped with the DMRs using BEDtools intersect to analyze how many peaks from tumor and normal organoids are located within the DMRs separately for H3K27ac ChIP-seq and ATAC-seq peaks.

#### Enrichment analysis

In order to detect possible enrichment of TFBS as well as ATAC-seq and ChIP-seq peaks in DMRs compared to the remaining genome, every DMR was matched with 1 million randomly picked genomic regions with similar length and CpG-count. The occurrence of the features of interest was then compared between DMRs and the average of the randomly chosen regions. The occurrence of the features of interest was then compared between DMRs and the average of the randomly chosen regions. A feature was defined as significantly enriched if its occurrence in the DMRs was larger than in the average of random regions in at least 95%, 99% or 99.9% of all comparisons (significance level < 0.05, < 0.01 or < 0.001).

### Targeted deep bisulfite sequencing

Targeted deep bisulfite sequencing was performed as described elsewhere(46). DMRs for validation were selected based on the following criteria: Proportion of hypermethylated DMRs reflects that of identified DMRs (> 90%), reversion in P12, conserved in humans, preferentially DMRs with at least one flanking transcript showing reverted expression in P12. Primer sequences are listed in Supplemental Table S1. The MiSeq (Illumina) run was conducted by the BioChip-Laboratory of the Essen University Hospital. Amplikyzer2 v. 1.2.0 was used for analysis. Due to the much higher coverage compared to WGBS, DMRs were classified as positively validated at a minimum methylation difference of 0.2.

### Whole genome sequencing

#### Sequencing and genome mapping

The Genomics and Proteomics Core Facility of the German Cancer Research Center (GPCF DKFZ, Heidelberg) performed the library preparation for WGS of parental and resistant cells of cell lines #3 and #9 as well as a control tail samples corresponding to line #3 with the TruSeq DNA PCR-free Kit (Illumina) and the 150 bp paired-end sequencing on a HiSeq X (Illumina).

Reads were aligned to the mouse reference genome GRCm38 using BWA-MEM 1. v. 0.7.15(43) with default settings. Duplicate reads were marked with sambamba v. 0.6.5(47).

#### Variant Calling

Variants and small InDels were called in a two-step process. First, candidate variants were called using freebayes v. 1.1.0(48) with parental and resistant tumor samples as well as a normal tail tissue.

In a second step, variants were validated and readjusted using Varlociraptor v. 1.1.1(49). The validated variants were separated into different groups according to their change in VAF between parental and resistant samples. Variants were defined as present in parental (VpPs) if they drop to a VAF of 0 from parental to resistant. Variants with a VAF > 0.1 in resistant and a VAF of 0 in parental were defined as present in resistant (VpRs). Variants with a VAF > 0 in both parental and resistant tumor samples were defined as present in parental and resistant (VpPRs).

VpPs and VpRs were validated in P5 and P12 tumors using the available WGBS data. Since the technical differences between WGBS and WGS affect the comparability of results from both methods, validation was performed solely on WGBS samples. VAFs in WGBS samples were called using Varlociraptor v. 1.1.1(49), with VpPs and VpRs from WGS as candidate variants. To adjust for bisulfite conversion, only A > T and T > A variants covered > 15x were kept for WGBS validation.

#### Data management and annotation

Snakemake v. 5.10(50) was used as workflow management system for the complete computational analysis.

Data management and visualization was performed using bcftools v. 1.9(51) and python v. 3.7 libraries seaborn v. 0.9 and pandas v. 0.24.

Variants were annotated using Jannovar v. 0.25(52) with the GRCm38 annotation database as well as SIFT scores (Ng and Henikoff, 2003) (Download source: http://sift.bii.a-star.edu.sg/sift4g/public//Mus_musculus/GRCm38.83.zip).

To compare SNV positions between both mouse lines, the closest variant positions between both variant calls were identified using BEDtools closest v. 2.27(38).

### Statistics

Replicates were performed as indicated in the figure legends. For statistical analyses R v. 3.6.0(53) and GraphPad Prism v. 7.03 were used. The applied test is described in the figure legends, respectively.

### Data Availability

Proteomics data have been deposited as a complete submission in the ProteomeXchange Consortium via the PRIDE partner repository (http://www.proteomexchange.org; data set identifier: PXD018093 and 10.6019/PXD018093). The .msf files obtained in Proteome Discoverer were converted into the mzIdentML standard format using ProCon PROteomics Conversion tool version 0.9.718 (PubMed ID 26182917). The RNA-seq generated during this study are available at GEO: GEO: GSE146348. The accession number for WGBS and WGS data deposited on ENA is: PRJEB37018.

### Code Availability

The custom code generated and used during the current study is available from the corresponding author on request.

## Results

### MEK_i_ resistance in PDAC is reversible upon drug withdrawal

To model MEK_i_ resistance in PDAC, we used primary low-passage cells derived from spontaneous PDAC of ten different *Ptf1a^wt/Cre^; Kras^wt/LSL-G12D^; Trp53^loxP/loxP^* mice, which develop aggressive and therapy-resistant tumors resembling key aspects of human PDAC(54, 55).

All primary cells lines (n=10) were sensitive to MEK_i_ with an IC50 in the low nanomolar range (5.44 nM to 41.91 nM, median 12.70 nM) (Supplemental Table S2). To induce MEK_i_ resistance, cells were treated with increasing trametinib doses over 3 to 4 months until they proliferated at 100-fold of the original IC50 dose (Fig. 1A). Resistance induction was successful in all 10 cell lines (Fig. 1B) and was accompanied by a strong block of ERK phosphorylation, underpinning the specificity of trametinib and suggesting a drug efflux independent resistance mechanism (Fig. 1C). To study the effect of drug withdrawal, one batch of resistant cells from each line was cultivated without MEK_i_ and samples were collected after 5 (P5) and 12 passages (P12) of drug withdrawal (Fig. 1A). Thereby a reversibility of the resistant phenotype was observed correlating with the duration of drug-free time (Fig. 1D).

**Fig. 1.**
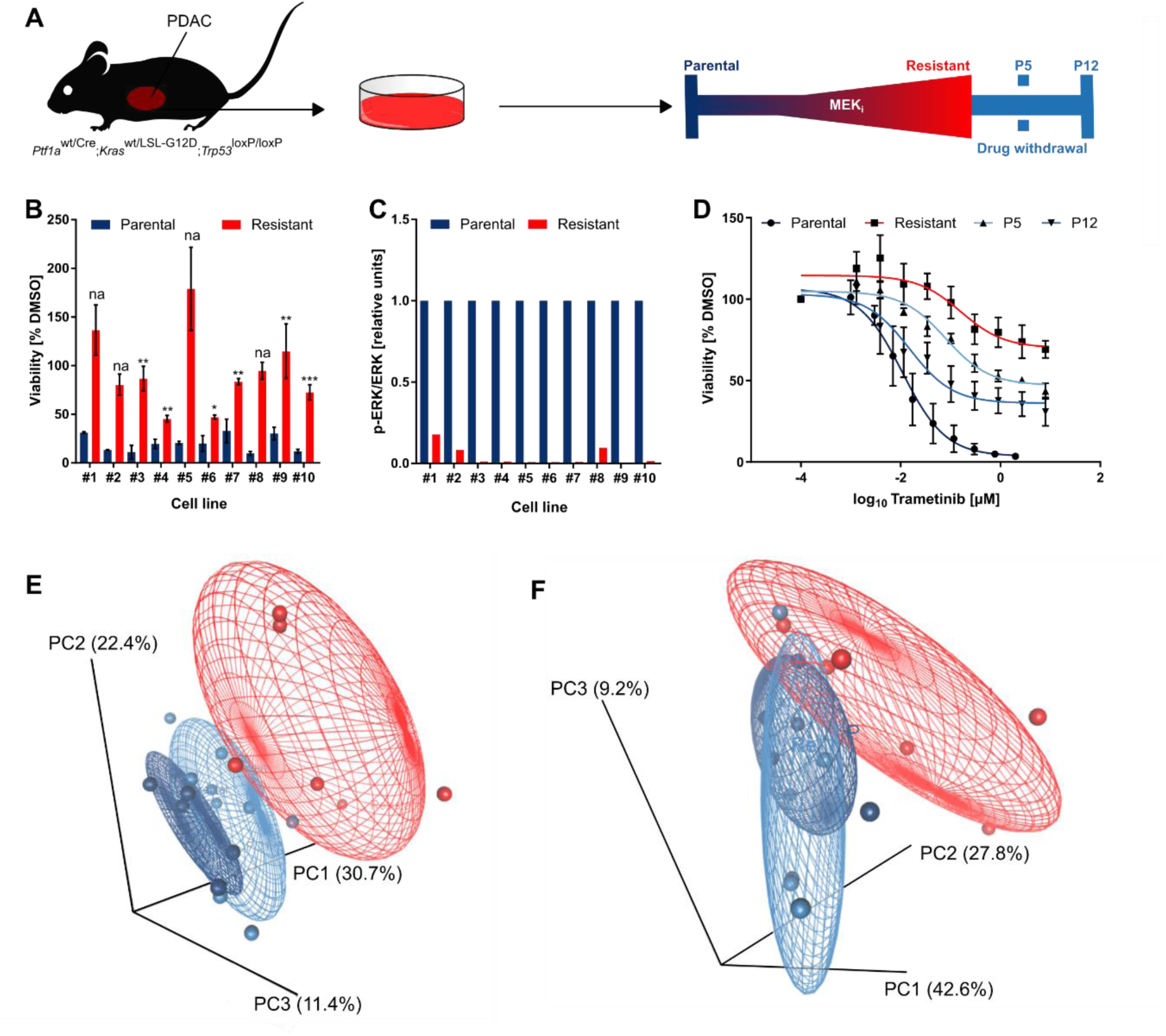
Acquired MEK_i_ resistance is reversible after drug withdrawal. **(A)** Timeline of resistance formation and drug withdrawal classified by different passages (P) without constant MEK_i_ treatment. **(B)** Bars represent the mean of two (na= two replicates in parental, three in resistant) or three independent cell viability measurements after 72 h in 300 nM MEK_i_ ± SD. Statistics was calculated by a two-tailed unpaired Student’s t-test on the log_2_ transformed DMSO-normalized values. **(C)** ERK phosphorylation in parental compared to MEK_i_-resistant cell states (p < 0.0001, two-tailed paired Student’s t-test on the log_2_ transformed ratios). **(D)** Resistance reversibility upon drug withdrawal. The mean of three independent experiments after 72 h incubation ± SD is shown for cell line #3 as representative example. **(E)** Principal component analysis of RNA-seq data between parental (dark blue), resistant (red) and reverting (P5 and P12; light blue) cell states. **(F)** Principal component analysis of all abundances identified by MS with more than one unique peptide.

We compared the expression profiles associated with acquired MEK_i_ resistance by performing RNA-seq of six matched parental, resistant and reverting cell states. Results of principal component analysis (PCA) over all expressed genes showed that the reverting states (P5 and P12) had a transcriptional profile more similar to the parental cells (Fig. 1E). The inter-individual variability explained considerably less of the transcriptional variation as opposed to the treatment. A similar sample separation could be observed on the protein level measured by mass spectrometry (Fig. 1F).

### Whole genome sequencing-based mutation analysis of MEK_i_-resistant cells

Genetic and non-genetic alterations may contribute to the development of a resistant phenotype(56). To determine whether genetic alterations were associated with MEK_i_ resistance, we performed whole genome sequencing (WGS) of two matched pairs of parental and resistant cell states with a median coverage of 40x. Compared to their treatment-naïve counterparts, resistant cells of lines #3 and #9 harbored 3657 and 3204 unique single nucleotide variants (SNVs), respectively. These variants are referred to as ‘variant present in resistant’ (VpR) (Fig. 2A,B, Supplemental Tables S4 and S5). A smaller number of variants was present in the parental, but could not be detected in the resistant cells (‘variant present in parental’, VpP) (131 in cell line #3 and 837 in cell line #9). Less than 1% of the VpRs were located in the coding regions (CDS) (36 in #3 and 23 in #9) (Supplemental Tables S4 and S5). Of these, 25 (cell line #3) and 19 (cell line #9) were nonsense or missense mutations, respectively (Supplemental Tables S4 and S5).

**Fig. 2.**
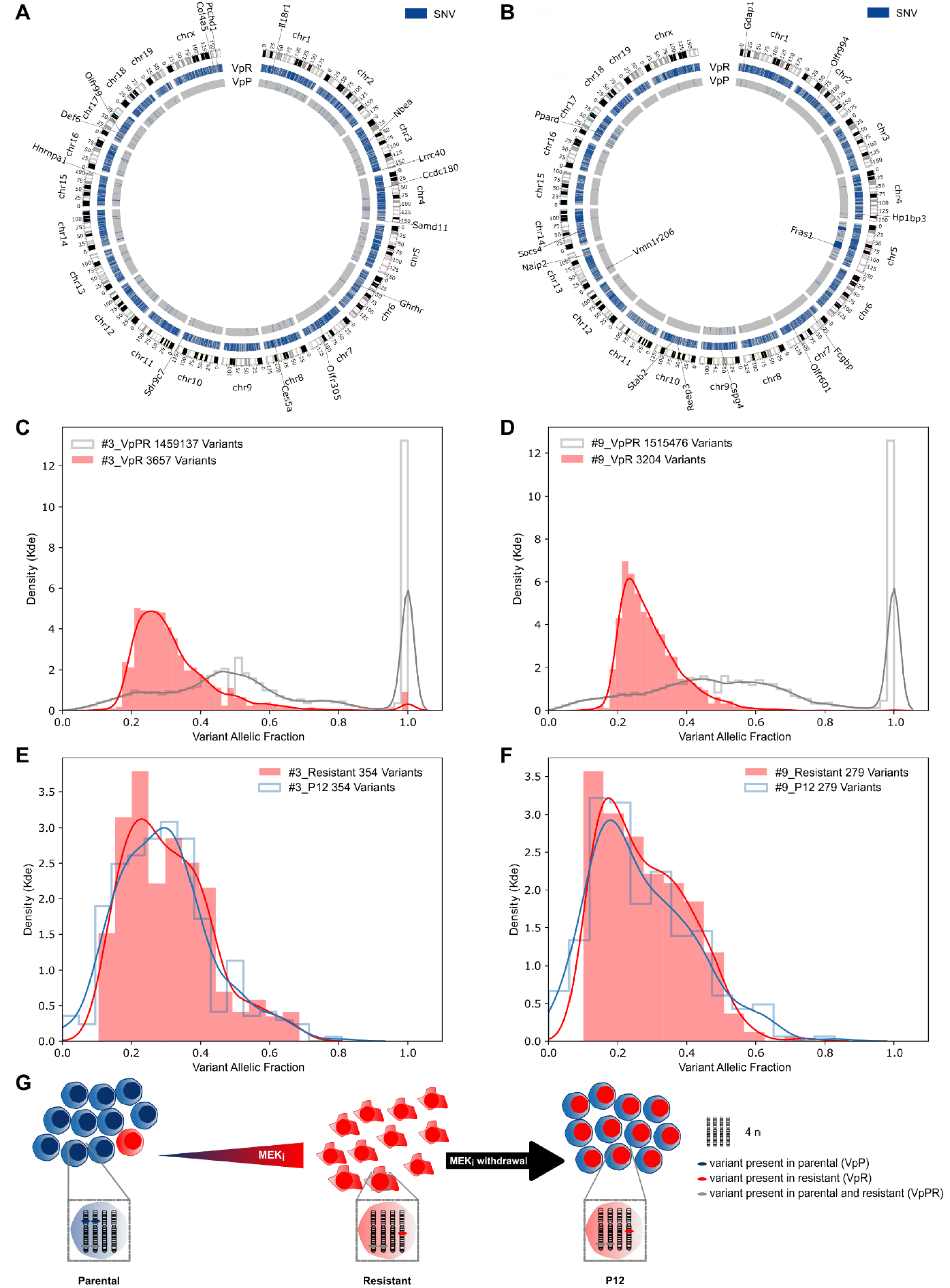
MEK_i_ resistance is based on clonal expansion, while reversion upon drug withdrawal is independent of a parental outgrowth. **(A,B)** Circos plots displaying VpRs or VpPs in cell lines #3 **(A)** and #9 **(B)**. Genes predicted as deleterious by SIFT are named. **(C,E)** Kernel density estimation (kde) for the VAF of VpRs in comparison to variants present in parental and resistant (VpPRs) for cell line #3 **(C)** or #9 **(D)**. **(E,F)** Density plot for the VAF of VpRs in resistant compared to P12 in cell line #3 **(E)** or #9 **(F)**. Only A > T and T > A variants called by WGS and validated by WGBS are shown. **(G)** Model of cell population dynamics during gain and loss of MEK_i_ resistance in PDAC.

SIFT prediction to assess the impact of these VpRs revealed 12 and 9 possibly deleterious variants for cell lines #3 and #9, respectively (Fig. 2A,B, Supplemental Tables S4 and S5). The putative variant effect supposed by Jannovar(52) was moderate for those classified as deleterious by SIFT. None of them were found in both cell lines or affected the same genes. None of the genes affected by the VpRs are known to be involved in the RAS-dependent MEK_i_ targeted MAPK or in the phosphoinositide 3-kinase (PI3K) pathway. In accordance, none of the mutant genes was listed in the COSMIC database to confer therapy resistance to human cancer cells. In addition to SNVs, small insertions and deletions (InDels) were called. However, we found none of the genes affected to be involved in re-activation or bypassing of the MEK pathway (Supplemental Tables S4 and S5). Overall, we did not identify any genetic variant that might explain the resistant phenotype.

### Whole genome sequencing reveals clonal expansion during acquired MEK_i_ resistance

We next addressed whether MEK_i_ resistance evolves by expansion of a subclone from the parental cells. Therefore, the distribution of the allele fraction of all variants found in resistant cells only (VpRs) was analyzed revealing a variant allele fraction (VAF) peak at around 0.25 in both cell lines (Fig. 2C,D). A potential explanation is the presence of this SNVs on one allele on a genetic background of four alleles e.g. in cells with tetraploidy as consequence of genome duplication. Polyploidization can be found in approximately 50% of human PDAC(57). By performing a cytogenetic analysis of metaphase chromosomes, we found in median 77 and 70 chromosomes per cell in parental cells of lines #3 and #9, respectively, which is in concordance with a near tetraploid karyotype (Supplemental Table S6). Evaluation of the VAF per chromosome revealed a lower chromosome count for chromosomes 9, 12 or 13 of cell line #3 or chromosomes 5 or 19 of #9 with VAF peak > 0.25, (Supplemental Fig. S1A,B). Taken together, the observed VAF distribution peak at 0.25 in these nearly tetraploid cells suggests that the cells with acquired MEK_i_ resistance are the result of clonal expansion of a single cell clone. Furthermore, most VpRs must have occurred after the incomplete genome duplications.

We next investigated whether a small proportion of parental cells that do not carry the VpRs might have survived the MEK_i_ treatment and then overgrown the VpR-containing resistant cells upon drug withdrawal. Therefore, we evaluated the VpR and VpR allele fractions in whole genome bisulfite sequencing (WGBS) data available for parental, resistant and reverting (P12) cells. Due to cytosine conversion by bisulfite modification, only a subset of 354 or 279 VpRs and 10 or 36 VpPs were available for further evaluation in cell lines #3 and #9, respectively. Nearly all VpRs were present in P12 cells with a VAF distribution similar to the resistant cells (Fig. 2E-G and Supplemental Fig. S1C,E). Furthermore, all but 2 VpPs, were absent in P12 cells (Supplemental Fig. S1D,F). Thus, despite rebounding MEK_i_ susceptibility, the resistant genotype persists in P12 cells.

Overall, MEK_i_ resistance is based on clonal expansion of a single cell clone, without evidence for mutations or structural variation in genes that could mediate MEK_i_-induced resistance. Loss of resistance upon drug withdrawal does not occur by outgrowth of parental cells.

### MEK_i_ resistance is associated with DNA hypermethylation

To address the potential involvement of epigenetic mechanisms in transcriptional variability, we applied a targeted drug screening approach addressing key epigenetic regulators including chromatin readers, histone modifiers and DNA methyltransferases in MEK_i_-resistant cells. We found a strong effect of the DNMT_i_ decitabine on the viability of resistant but not parental cells (Fig. 3A-C). The difference was by far more pronounced compared to inhibitors of other epigenetic regulators such as bromodomain and extracellular terminal (BET) proteins, class I-specific or pan-histone deacetylases (HDAC), suggesting that MEKi-induced formation of resistance involves critical changes in DNA methylation (Fig. 3A,B).

**Fig. 3.**
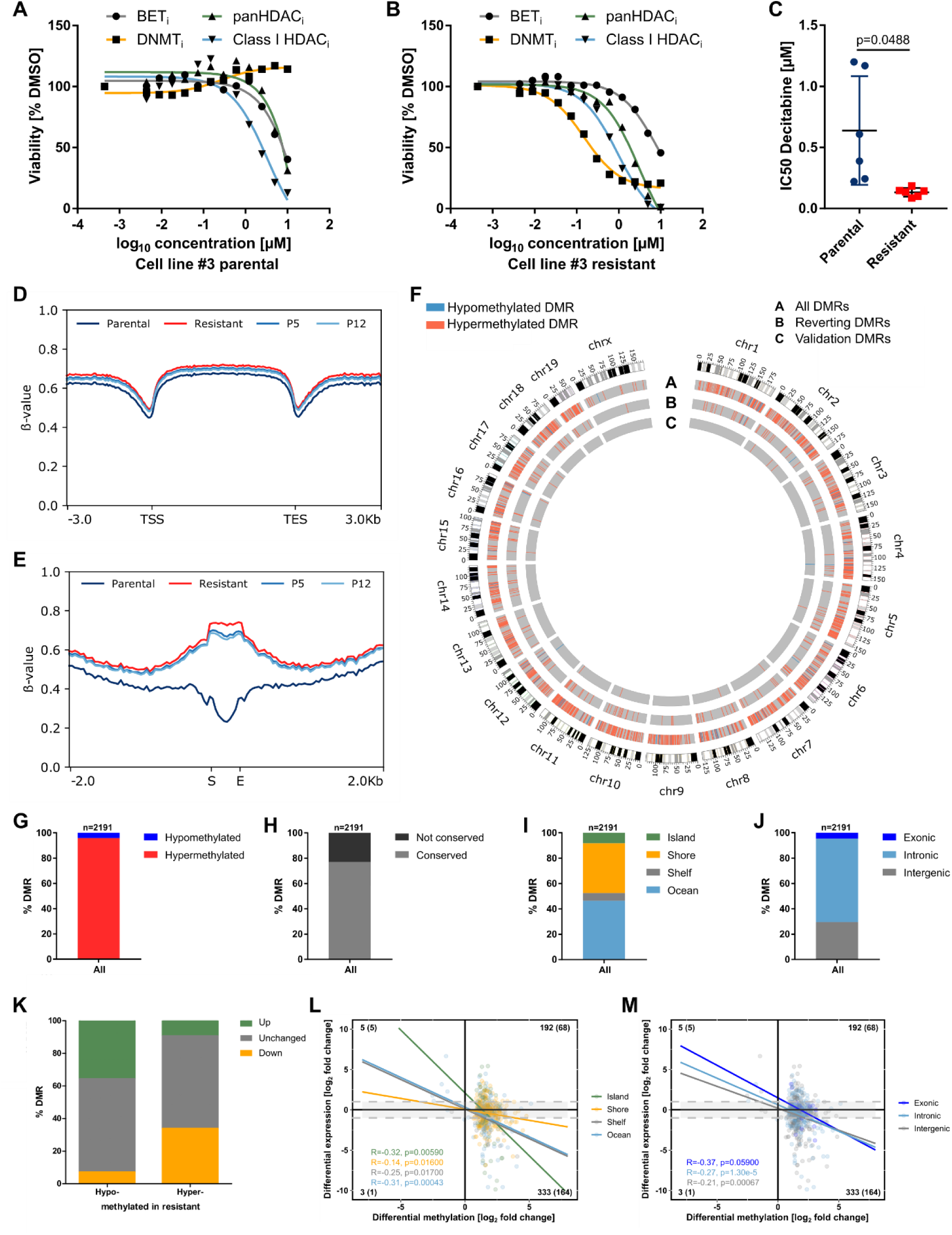
Whole genome bisulfite sequencing revealed DMRs mainly hypermethylated in MEK_i_ resistant cells. **(A,B)** Cell viability screen using inhibitors that target different epigenetic mechanisms in parental **(A)** or resistant cells **(B)**. Dose response curves for cell line #3 determined after 72 h using the CellTiter-Glo® cell viability assay are shown. **(C)** The IC50 of DNMT_i_ was significantly lower in six matched-pairs of parental versus resistant cells (two-tailed paired Student’s t-test). **(D)** Mean gene body methylation levels in parental, resistant, P5 and P12 cell states of cell lines #3 and #9. **(E)** Mean methylation levels of all DMRs and their flanking regions (± 2 kb) in the four indicated cell states of two different cell lines (#3, #9). S indicates the DMR start and E its end. **(F)** Circos plot indicates the chromosomal location of 2191 DMRs between two parental and resistant cell lines (circle A). A scoring function was developed to define 217 reverting DMRs whose methylation pattern in P12 resembles that of parental cells (circle B). Circle C displays 15 DMRs that were validated by targeted deep bisulfite sequencing. **(G)** Proportion of DMRs hypo- or hypermethylated in resistant cells. **(H)** Degree of mouse-human DMR-sequence conservation according to the UCSC liftover tool. **(I,J)** Relative location of DMRs in relation to CpG islands **(I)** or genes **(J)**.

To assess MEK_i_ treatment-induced alterations in genome-wide DNA methylation, we performed whole genome bisulfite sequencing of two cell lines, each at four different states: parental, resistant and reverting cells at P5 and P12. Overall gene body and promotor methylation levels between all four cell states remained unchanged (Fig. 3D). We defined differentially methylated regions (DMRs) with a minimum CpG-count of four and a minimal difference in methylation level of 0.4. Thereby, 2191 DMRs relatively equally distributed over all chromosomes were found when comparing parental and resistant cells (Fig. 3E,F and Supplemental Table S7). These DMRs covered a total of 38,031 CpGs with a mean CpG content per DMR of 17 (min=4, max=178) and an average length of 794 bp (min=12 bp, max=4157 bp) (Supplemental Table S7). Remarkably, more than 96% of these DMRs were hypermethylated in the resistant cells (Fig. 3F,G). The nucleotide sequence of the majority of 1756 DMRs was conserved in humans based on the UCSC liftover tool corresponding to a degree of conservation > 76% (Fig. 3H and Supplemental Table S7). 43.37% were located in ocean areas of the genome, while 39.25% and 6.12% were present in the CpG island flanking shores and shelves, respectively. Only 8.26% of all DMRs overlapped with a CpG island (Fig. 3I). Approximately one third (29.53%) of the DMRs was located in intergenic regions, while the others were present in intragenic, predominantly intronic regions (65.95%) (Fig. 3J).

We next addressed whether the DMR methylation is correlated with the expression of neighboring genes. We annotated two flanking genes to each DMR based on the nearest transcription start site (TSS) in each direction. Gene expression was correlated using RNA-seq data of the matching cell line (Fig. 3K and Supplemental Table S7). More than half (61%) of the flanking genes were not differentially expressed between parental and resistant cells. One-third of genes flanking hypomethylated DMRs was upregulated and about 25% of genes associated with hypermethylated DMRs were downregulated. Limiting the analysis to genes located downstream of each DMR, the overall inverse correlation remained poor independent of the DMR position in the genome (Supplemental Fig. S2A,B). Focusing only on genes whose TSS was located less than 7.5 kb downstream of a DMR resulted in a higher negative correlation of DNA methylation and transcript level for DMRs associated with island, ocean as well as intragenic regions (Fig. 3L,M). In summary, we identified extensive hypermethylation following acquired MEK_i_ resistance, supporting epigenetic plasticity in our model system.

### A DMR subset reverts upon MEK_i_ withdrawal

We next addressed the question if DMRs involved in MEK_i_ resistance may revert upon drug withdrawal (Fig. 4A). We identified 217 DMRs, from here on referred to as reverting DMRs, whose methylation status correlated with MEK_i_ sensitivity at all stages analyzed (Fig. 4B,C and Supplemental Table S7). This correlation was further supported by the observation that the degree of methylation in P5 cells with intermediate resistance phenotype was always between the value of the resistant cells and the P12 cells (Supplemental Fig. S2C,D).

**Fig. 4.**
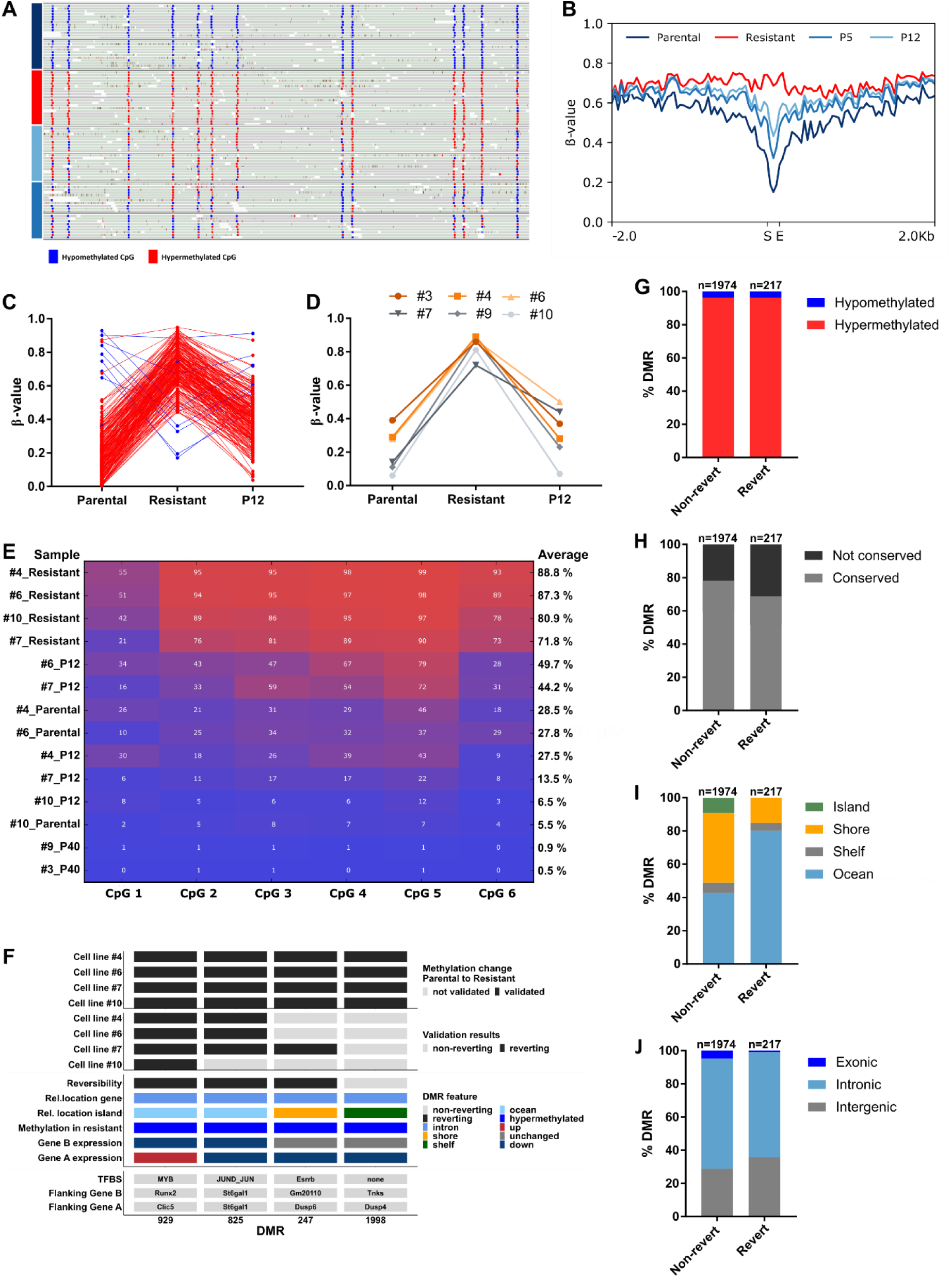
Distinct DMRs revert after MEK_i_ withdrawal. **(A)** Representative Integrative Genomics Viewer snapshot of reverting DMR_1716 in the parental (dark blue), resistant (red), P5 (light blue) and P12 (blue) cell states of cell lines #3 and #9 (upper row and lower row, respectively). 13 of 16 CpGs are shown. **(B)** Mean methylation levels of the reverting DMRs and their 2 kb up- and downstream regions of cell lines #3 and #9 in the parental, resistant, P5 and P12 cells (S: Start DMR; E: End DMR). **(C)** Methylation pattern of 217 reverting DMRs divided into hypo- (blue) and hypermethylated (red) regions. **(D,E)** Methylation pattern of DMR_929 in four independent cell lines compared to the WGBS samples based on the average methylation b-value in the region **(D)** or CpG wise **(E)**. In addition, P40 samples of lines #3 and #9 previously analyzed by WGBS were included in the validation. **(F)** Example of 4/15 DMRs validated by targeted deep bisulfite sequencing in four independent cell lines. Non-reverting DMR_1998 served as a negative control. **(G-J)** Comparison of reverting and non-reverting DMRs based on methylation change in resistant cells **(G)**, sequence conservation in human **(H)** and their relative location to CpG islands **(I)** or genes **(J)**.

We validated a set of 15 selected DMRs in four additional cell lines in parental, resistant and P12 states using targeted deep amplicon bisulfite sequencing(46). Ten out of 15 DMRs were differentially methylated between parental and resistant cells in at least two of four cell lines used for validation. In addition, eight of these 10 DMRs showed a reverting DNA methylation in the P12 state (Fig. 4D-F and Supplemental Fig. S2E and S3). In particular, the DNA methylation levels of reverting DMRs in cell lines #3 and #9 remained at P12 levels or below even after 40 passages under MEK_i_ withdrawal (Fig. 4E and Supplemental Fig. S3).

Comparing various features of reverting and non-reverting DMRs, we found the proportion of hypomethylated DMRs to be the same in both groups (Fig. 4G). The number of human-mouse conserved DMRs was slightly lower in the reverting DMRs (Fig. 4H). Reverting DMRs were more frequently located in the ocean than non-reverting DMRs (80.18% vs. 42.65% in non-reverting DMRs), while they were underrepresented in shores (Fig. 4I). Notably, none of the reverting DMRs overlapped with a CpG island. In exonic regions, reverting DMRs were less frequent (0.92%) compared to non-reverting DMRs (4.91%) (Fig. 4J).

### *In silico* evaluation of regulatory relevance of reverting DMRs

To investigate the potential relevance of the reverting DMRs, we analyzed their co-localization with miRNA target regions, VISTA enhancers and transcription factor binding sites (TFBS) as annotated by Ensembl. (Fig. 5A and Supplemental Table S7). Compared to randomly selected regions with similar length and CpG content, only TFBS were significantly enriched in all 2191 DMRs and even more pronounced in the 217 reverting DMRs (Fig. 5B).

**Fig. 5.**
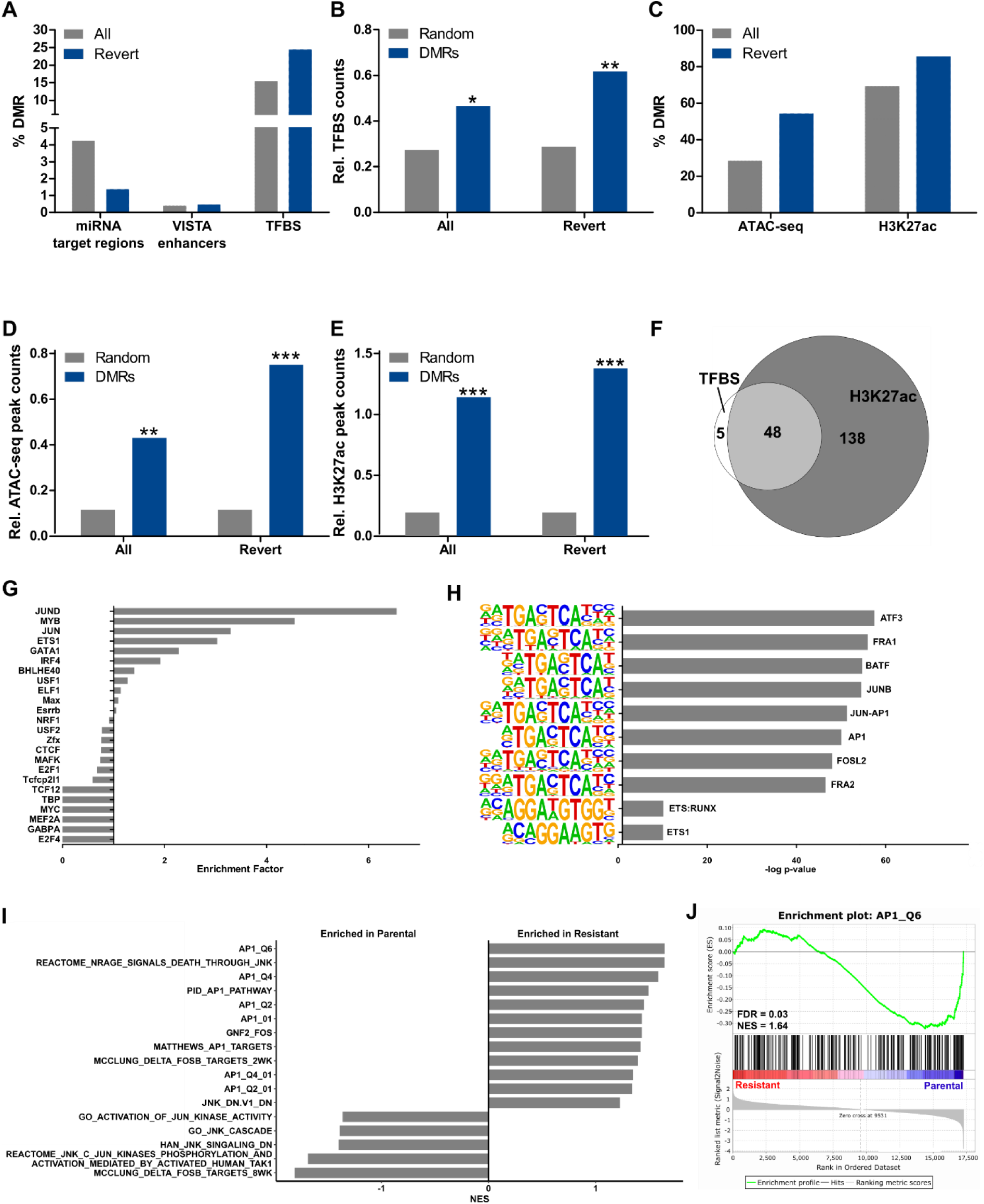
Functional relevance of DMRs and DNA methylation in MEK_i_ resistance. **(A)** Percent of DMRs overlapping with the indicated feature. **(B)** TFBS were significantly enriched in all as well as in reverting DMRs compared to 10^6^ random regions of similar length and CpG count (** significance level < 0.01; * significance level < 0.05). **(C)** Percent of DMRs that overlap with an ATAC-seq or ChIP-seq H3K27ac peak found in at least 2 organoids by Roe and co-workers (41). **(D,E)** Enrichment for ATAC-seq **(D)** or H3K27ac **(E)** peaks relative to 10^6^ random regions of similar length and CpG count (*** significance level < 0.001; ** significance level < 0.01). **(F)** Venn diagram of reverting DMRs’ overlap with TFBS and/or H3K27ac peaks. **(G)** Enrichment of TFBS for the indicated transcription factors in the 217 reverting DMRs compared to 1974 non-reverting DMRs. **(H)** Enrichment of TFBS motifs from the homer database between the reverting DMRs and random background sequences. **(I)** NES of AP1 related gene expression signatures that are significantly (FDR < 0.25) different between parental and resistant cells based on GSEA of RNA-seq data. **(J)** Enrichment plot of the AP1_Q6 gene expression signature.

The activity of regulatory elements is highly tissue- as well as context-specific and their deregulation is often observed in cancer. Therefore, we aligned the identified DMRs with sequences reported to represent open chromatin (assay for transposase-accessible chromatin and sequencing (ATAC-seq)) or potential active enhancer sites (chromatin immunoprecipitation and sequencing (ChIP-seq) for H3K27ac) in murine pancreas cells(41). These data were obtained from murine organoids, similar to our model, derived from PDAC of KPC mice (*Kras^wt/LSL-G12D^; Trp53^wt/LSL-R172H^; Pdx1-Cre*). We found that 28.5% of all 2191 DMRs overlapped with ATAC-seq peaks, which comprise only about 0.8% of the genome, while 3% of the genome, but 69.3% of all DMRs overlapped with H3K27ac ChIP-seq peaks (Fig. 5C and Supplemental Table S7). Thus, DMRs were significantly enriched for open and/or active (H3K27ac histone occupied) chromatin regions compared to random regions of similar length and CpG content (Fig. 5D,E). In both cases, this enrichment was even more pronounced for reverting DMRs. More than 90% of TFBS-containing reverting DMRs overlapped with a H3K27ac-marked region, which underlines their potential regulatory relevance in MEK_i_-resistant PDAC cells (Fig. 5F).

Using two independent enrichment analysis tools, we found that binding motifs for proteins belonging to the activator protein 1 (AP1) family were amongst the top enriched TFBS (Fig. 5G,H). It is well documented that AP1 binding to its respective motif is strongly dependent on the methylation status of its recognition site and proximity(58, 59). AP1 is a protein dimer formed by members of the JUN, FOS and ATF families. Notably, JUN was among the proteins regulated in MEK_i_-treated PDAC cells. Resistant cells displayed both an increased total protein expression and enhanced JUN phosphorylation at Ser73 (Supplemental Fig. S4). Furthermore, different AP1/JUN expression signatures were enriched in the parental cell lines (Fig. 5I,J).

In conclusion, MEK_i_ resistance is associated with DNA hypermethylation at regulatory elements including TFBS and active enhancer sites located in open chromatin. We identified the AP1 transcription factor complex as a potentially crucial factor in mediating MEK_i_-induced resistance.

### CASP3 down-regulation counteracts treatment-induced apoptosis in MEK_i_-resistant cells

The AP1 transcription factor family is involved in apoptosis where CASP3 activation is the final step in the apoptosis signaling cascade. Two of the DMRs whose methylation correlates with the MEK_i_ sensitivity were located in close proximity downstream of the *Casp3* locus (DMR_2004, DMR_2005; Fig. 6A and Supplemental Table S7). Inversely correlating with DMR hypermethylation, *Casp3* mRNA and CASP3 protein expression were significantly reduced in MEK_i_-resistant cells compared to their naïve counterparts, while reaching their original level in reverting P12 cells (Fig. 6B,C).

**Fig. 6.**
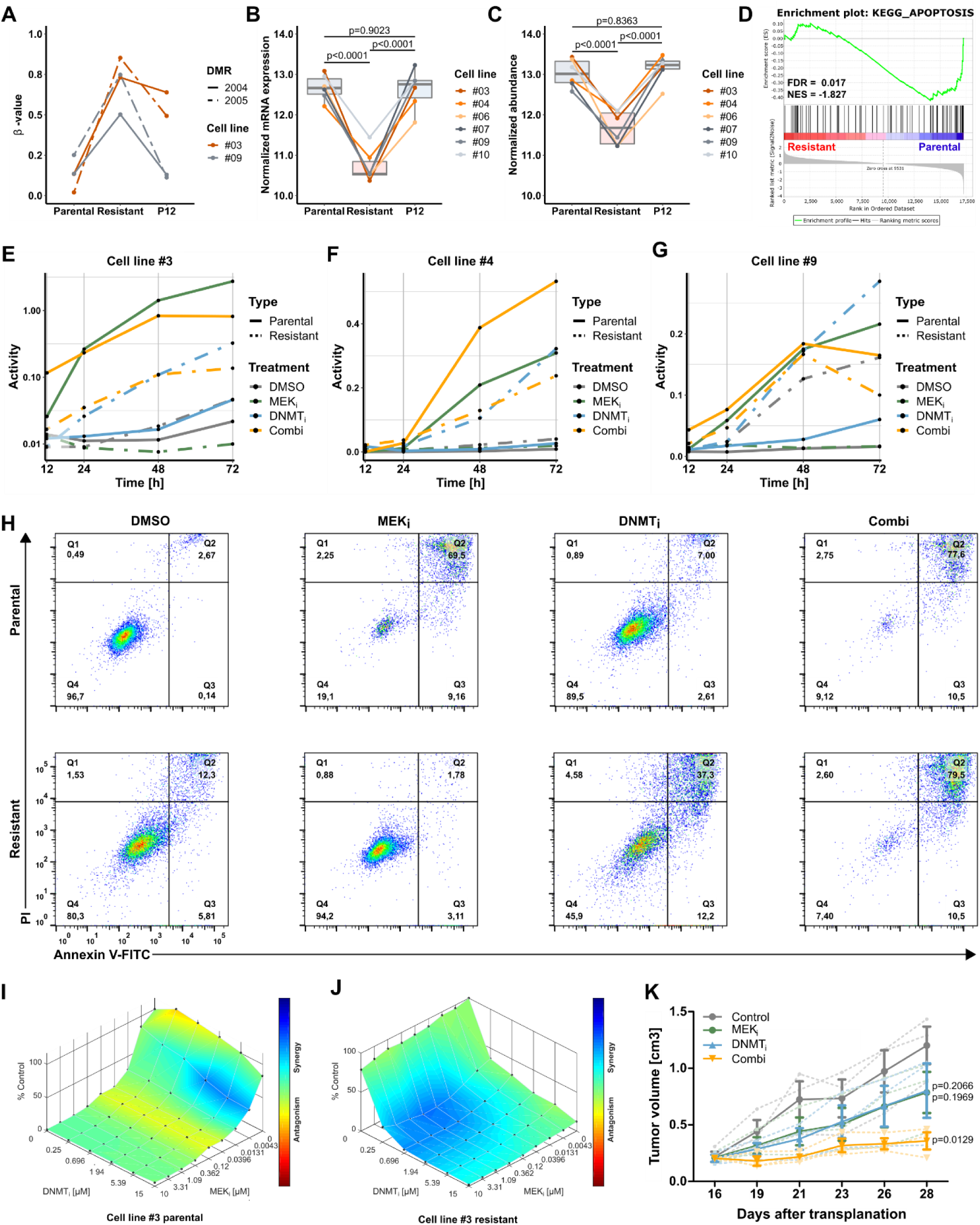
Adaptive CASP3 regulation and apoptosis during gain and loss of MEK_i_ resistance. **(A)** Methylation pattern of DMR_2004 and DMR_2005 that were hypermethylated in resistant cells and reverted upon MEK_i_ withdrawal in P12. **(B)** *Casp3* mRNA expression measured by RNA-seq in six matched pairs of parental, resistant and P12 cells. The adjusted p-values based on a differential gene expression analysis using DESeq2 are shown. **(C)** Mass spectrometry revealed a reverting CASP3 protein expression in six matched pairs of parental, resistant and P12 cells. Significance was determined by ANOVA and Tukey post-hoc test. **(D)** GSEA of RNA-seq data revealed a significantly enriched KEGG pathway apoptosis gene set in parental compared to resistant cells. **(E-G)** CASP3 activity assay after incubation with the indicated compounds. **(H)** FACS analysis of Annexin V-FITC/PI co-stained parental (upper row) or resistant cells (lower row) of line #9 upon 84 h of indicated treatment. **(I,J)** Synergy analysis of MEK_i_ plus DNMT_i_ using the Loewe method of the Combenefit software shown for cell line #3 parental **(I)** and resistant **(J)**. **(K)** Growth curves of PDX treated with MEK_i_, DNMT_i_ or the combination. Solid lines represent the mean tumor volume of three mice per treatment group ± SEM. Statistical significance versus control was determined using a two-tailed unpaired Student’s t-test.

Functional annotations revealed an overlap of both DMR_2004 and DMR_2005 with active enhancers (H3K27ac marked) and open chromatin suggesting their relevance for *Casp3* transcriptional regulation (Supplemental Table S7). Furthermore, Gene Set Enrichment Analysis (GSEA) of KEGG pathways in the parental cells showing *Casp3* expression revealed an enrichment for genes involved in apoptosis compared to resistant cells with decreased CASP3 expression levels (FDR < 0.05, NES=-1.827) (Fig. 6D).

Notably, CASP3 activity was induced in the parental cell lines upon MEK_i_ treatment but down-regulated in the MEK_i_-resistant cells (Fig. 6E-G). In contrast, MEK_i_ withdrawal was accompanied by an increasing basal CASP3 activity in the resistant cells, which could be further enhanced by DNMT_i_ treatment. The DNMT_i_ effect on the CASP3 activity of the parental cells was only marginal, which is in accordance with the previously shown cell viability data. A combined treatment with MEK_i_ and DNMT_i_ induced the activity of CASP3 in MEK_i_-resistant cells similar as DNMT_i_ alone, whereas the impact on the parental cells was comparable to MEK_i_ treatment alone (cell lines #3 and #9, Fig. 6E,G) or even stronger (cell line #4, Fig. 6F). Finally, FACS analysis using an Annexin V/propidium iodide co-staining confirmed the suppression of apoptosis by CASP3 inactivation as a potential MEK_i_ resistance mechanism in PDAC (Fig. 6H and Supplemental Fig. S5).

To evaluate the synergism between MEK_i_ and DNMT_i_ we performed *in vitro* cell viability assays. In resistant cells, a synergistic effect of MEK_i_ and DNMT_i_ was observed even at low doses of DNMT_i_. Synergism in the parental cells was only observed at high DNMT_i_ concentrations (Fig. 6I,J and Supplemental Fig. S6A-F).

Our data revealed that MEK_i_ resistance was accompanied by a downregulation of CASP3 together with a loss of activity which recovered after drug withdrawal. MEK_i_-induced apoptosis was impaired in the resistant cells but could be overcome by concomitant DNMT_i_ treatment. In addition, we investigated the effect of MEK_i_ and DNMT_i_ inhibition using a human *in vivo* setting. We treated three different PDX models with either MEK_i_, DNMT_i_ or a combination of both and observed a synergistic effect in all three models (Fig. 6K and Supplemental Fig. S6G,H). Overall, our data show a key relevance of DNA methylation for maintaining MEK_i_ resistance in PDAC, which results in a remarkable DNMT_i_ vulnerability.

## Discussion

PDACs are characterized by a remarkable resistance to virtually all therapeutic strategies. Several aspects challenge the unravelling of underlying mechanisms(60). A key limitation of studies in advanced PDAC is the lack of longitudinal sample acquisition in individual treated patients due to anatomical, ethical and logistical reasons among others.

Here, we utilized an *in vitro* model based on low-passage primary cells derived from tumors of a genetically engineered PDAC mouse model(54) to analyze molecular changes that arise under MEK inhibition. The use of tumor cell populations with the same genetic background minimizes the influence of confounding factors such as inter-individual heterogeneity or contaminating non-tumor cells, the latter being a major concern when using primary PDAC tumor tissue. Consequently, our model cannot account for non-tumor cell-intrinsic mechanisms such as involvement of the tumor microenvironment(61). However, the observation that we were able to induce MEK_i_ resistance in all ten cell lines suggests that, at least in our model system, resistance formation is mediated by tumor cell intrinsic mechanisms.

Using WGS, we clearly demonstrate that the resistant cell clones evolve from a single precursor cell in the parental cell population. It is not possible to distinguish whether the genetic variants observed in the resistant cells were already present in the parental cells or arose during MEK_i_ treatment or a combination of both. The observed clonality strongly indicates that the parental cell population is composed of cells with different abilities to adapt to MEK_i_ treatment with single cells having the potential to develop a resistance phenotype.

In a human PC-9 lung cancer model Hata et al.(62) compared gefitinib resistant cells harboring a resistance mediating *EGFR* mutation with resistant cells that expanded under treatment without such mutations. Gefitinib-resistant *EGFR* mutant PC-9 cells had a similar transcriptional profile to their parental gefitinib-naive cell pool(62). In contrast, transcriptomic differences found between our mouse parental and MEK_i_-resistant cells were also described for PC-9 cells without *EGFR* resistance mutation(62). Furthermore, it took about four month from the onset of drug exposure to a fully MEK_i_-resistant cell population(62). This is a similar time frame to that observed for the PC-9 lung cells without a known resistance-mediating *EGRF* mutation(62). Whereas PC-9 cell pools harboring an *EGFR* resistance mediating mutation, developed resistance within 6 weeks(62). Given the similar proliferation rates of cells in our PDAC mouse model and the PC-9 cells this argues against a mutation in a classical resistance gene as underlying cause of MEK_i_ resistance. Consistently, using WGS we did not detect mutations in genes involved in re-activation or bypassing of the targeted MAPK pathway in the resistant cells. Although this approach cannot definitively rule out the presence of such mutations, it provides clues for the presence of alternative mechanisms that confer MEK_i_ resistance to cells. This is further supported by the observation that cells lose their resistance phenotype during drug withdrawal without re-gaining the parental genotype. As it is unlikely that the formation of resistance is a direct consequence of mutations in genes involved in drug resistance, we hypothesize that the predisposition of the originating cell could be due to an epigenetic plasticity that enables it to adapt better to the environmental conditions than other cells e.g. by DNA methylation changes. Whether this plasticity is the consequence of sequence alterations already present in the parental cell or due to stochastic epigenetic variation remains to be determined. An example for an epigenetic factor contributing to phenotypic adaptation upon drug treatment is described in a study by Wang et al.(21) showing that modulation of histone methylation is involved in MEK_i_ resistance.

As DNA methylation is an important epigenetic mediator already known to be involved in the progression of PDAC or other tumors, as well as the formation of distant metastases, we determined DNA methylation changes in our PDAC model(63). To our knowledge, we provide the first WGBS dataset of therapy-resistant PDAC, albeit in murine cells, which provides an unbiased and comprehensive view at the dynamic changes of the methylome in cells in response to drug exposure and subsequent drug withdrawal. The differentially methylated regions showed a high degree of conservation between the mouse and human genome and overlapped with known regulatory elements like TFBS and potential enhancer sites, which supports their functional relevance in gene regulation. We could also identify a subset of reverting DMRs whose gain and loss of methylation reflected gain and loss of the resistant phenotype. Among these reverting DMRs, binding sites for the dimeric transcription factor complex AP1 were significantly enriched. Its DNA hypermethylation, as present in the resistant cell states, is known to prevent AP1 binding(58, 59).

We identified a potential *Casp3* enhancer, whose methylation status correlated inversely with CASP3 expression and was dependent on MEK_i_-presence in the culture medium. A functional role of the methylation is supported by the results from DNMT_i_ treatment which results in CASP3 re-activation in resistant cell lines. Deregulation of CASP3 is a common mechanism for tumor cells to mediate therapy resistance(64–68). Here, we show the association of CASP3 activity and methylation of distal DNA regions related to modulation of treatment-induced apoptosis.

It is a matter of debate if DNA methylation changes are causally involved in the regulation of gene expression or if they develop downstream of transcription factor-mediated gene regulation(69). Interestingly, the DNA methylation changes that occur during drug exposure and resistance formation are almost exclusively DNA hypermethylation events. Such *de novo* methylation of previously unmethylated CpGs depends on the activity of methyltransferases DNMT3A und DNMT3B. Therefore, blocking *de novo* methylation could prevent or impede the formation of MEK_i_ resistance in PDAC cells or primary tumors. Indeed, the synergistic effect of combined MEK_i_ and DNMT_i_ treatment observed in resistant PDAC cells strongly suggest that MEK_i_ resistance is attenuated by decitabine treatment. Decitabine is an inhibitor that blocks the activity of both *de novo* methyltransferases and of the maintenance methyltransferase DNMT1. It will be interesting to evaluate if drug resistance in general is associated with DNA hypermethylation or if this observation is restricted to MEK inhibitors in PDAC cells.

Overall, our results of a MEK_i_ adaptive DNA hypermethylation landscape in a single cell clone support epigenetic plasticity of tumor cells as a driver in PDAC therapy resistance. The remarkable DNMT_i_ sensitivity might inspire new combinatory therapeutic approaches to overcome therapy resistance in PDAC.

## Acknowledgments

We thank Kristin Fuchs (Medizinisches Proteom-Center, Bochum) for the performance of mass spectrometry; Elke Jürgens and Regina Kubica (Department of Human Genetics, Essen) for cytogenetics. Furthermore, we thank the High Throughput Sequencing Unit (Genomics & Proteomics Core Facility, DKFZ) for providing excellent whole genome and whole genome bisulfite sequencing services and the CeGaT GmbH (Tübingen) for RNA-sequencing. We thank Ludger Klein-Hitpass (Biochip-Labor, University Hospital Essen) for providing excellent amplicon-sequencing services. We thank Fung-Yi Cheung and Smiths S. Lueong (both Institute for Developmental Cancer Therapeutics, University Hospital Essen) for technical advice and discussions. We thank Florian Schmidt (Max-Planck-Institut für Informatik, Saarbrücken) for helpful discussions regarding the integration of RNA-seq and WGBS data.

J.T.S is supported by the German Cancer Consortium (DKTK), the Deutsche Forschungsgemeinschaft (DFG) through grant SI1549/3-1 (Clinical Research Unit KFO337) and SI1549/4-1; the Deutsche Krebshilfe (German Cancer Aid) through #70112505, PIPAC and #70113834, PREDICT-PACA; the European Uniońs Seventh Framework Programme for research, technological development and demonstration (FP7/CAM-PaC) under grant agreement no° 602783. K.U.L is supported by the DFG (LU-1944/3-1). A.S and R.T.L. are supported by Associazione Italiana Ricerca sul Cancro (AIRC 5×1000 n. 12182) and Fondazione Cariverona: Oncology Biobank Project “Antonio Schiavi” (prot. 203885/2017). This work was supported by PURE (Protein research Unit Ruhr within Europe) funded by the Ministry of Science, North Rhine-Westphalia, Germany.

## Author Contributions

Conceptualization, L.K.G., M.Z., J.T.S.

Methodology, L.K.G., J.F., S.T.L., C.S., J.K., L.H., K.U.L., M.T-A., D.B., A.S., R.T.L., K.E.W., B.S., M.Z.

Software, L.K.G., J.F., S.T.L., C.S., J.K., L.H., K.U.L., S.R.

Formal Analysis, L.K.G., J.F., S.T.L., C.S., J.K., L.H., K.U.L., K.E.W.

Resources, D.B., A.S., R.T.L., J.T.S.

Writing – Original Draft, L.K.G., J.F., M.Z., J.T.S.

Writing – Review & Editing, S.T.L., B.S., S.A.J., S.R., B.H., M.Z., J.T.S.

Visualization, L.K.G., J.F.

Supervision, S.T.L., S.R., B.H., M.Z., J.T.S.

Funding Acquisition, J.T.S.

## References

1. Cancer Genome Atlas Research Network. Integrated genomic characterization of pancreatic ductal adenocarcinoma. Cancer Cell 2017;32:185–203

2. Waddell N, Pajic M, Patch AM, Chang DK, Kassahn KS, Bailey P, et al. Whole genomes redefine the mutational landscape of pancreatic cancer. Nature 2015;518:495–501

3. Chan-Seng-Yue M, Kim JC, Wilson GW, Ng K, Figueroa EF, O’Kane GM, et al. Transcription phenotypes of pancreatic cancer are driven by genomic events during tumor evolution. Nat Genet 2020;52:231–40

4. Ohren JF, Chen H, Pavlovsky A, Whitehead C, Zhang E, Kuffa P, et al. Structures of human MAP kinase kinase 1 (MEK1) and MEK2 describe novel noncompetitive kinase inhibition. Nat Struct Mol Biol 2004;11:1192–7

5. Collisson EA, Trejo CL, Silva JM, Gu S, Korkola JE, Heiser LM, et al. A central role for RAF-->MEK-->ERK signaling in the genesis of pancreatic ductal adenocarcinoma. Cancer Discov 2012;2:685–93

6. Walters DM, Lindberg JM, Adair SJ, Newhook TE, Cowan CR, Stokes JB, et al. Inhibition of the growth of patient-derived pancreatic cancer xenografts with the MEK inhibitor trametinib is augmented by combined treatment with the epidermal growth factor receptor/HER2 inhibitor lapatinib. Neoplasia 2013;15:143–55

7. Vena F, Li Causi E, Rodriguez-Justo M, Goodstal S, Hagemann T, Hartley JA, et al. The MEK1/2 inhibitor pimasertib enhances gemcitabine efficacy in pancreatic cancer models by altering ribonucleotide reductase subunit-1 (RRM1). Clin Cancer Res 2015;21:5563–77

8. Van Laethem JL, Riess H, Jassem J, Haas M, Martens UM, Weekes C, et al. Phase I/II study of refametinib (BAY 86-9766) in combination with gemcitabine in advanced pancreatic cancer. Target Oncol 2017;12:97–109

9. Infante JR, Somer BG, Park JO, Li C-P, Scheulen ME, Kasubhai SM, et al. A randomised, double-blind, placebo-controlled trial of trametinib, an oral MEK inhibitor, in combination with gemcitabine for patients with untreated metastatic adenocarcinoma of the pancreas. Eur J Cancer 2014;50:2072–81

10. Chung V, McDonough S, Philip PA, Cardin D, Wang-Gillam A, Hui L, et al. Effect of selumetinib and MK-2206 vs axaliplatin and fluorouracil in patients with metastatic pancreatic cancer after prior therapy: SWOG S1115 study randomized clinical trial. JAMA Oncol 2017;3:516–22

11. Van Cutsem E, Hidalgo M, Canon JL, Macarulla T, Bazin I, Poddubskaya E, et al. Phase I/II trial of pimasertib plus gemcitabine in patients with metastatic pancreatic cancer. Int J Cancer 2018;143:2053–64

12. Fedele C, Ran H, Diskin B, Wei W, Jen J, Geer MJ, et al. SHP2 inhibition prevents adaptive resistance to MEK inhibitors in multiple cancer models. Cancer Discov 2018;8:1237–49

13. Ruess DA, Heynen GJ, Ciecielski KJ, Ai J, Berninger A, Kabacaoglu D, et al. Mutant KRAS-driven cancers depend on PTPN11/SHP2 phosphatase. Nat Med 2018;24:954–60

14. Lin L, Sabnis AJ, Chan E, Olivas V, Cade L, Pazarentzos E, et al. The Hippo effector YAP promotes resistance to RAF- and MEK-targeted cancer therapies. Nat Genet 2015;47:250–6

15. Ponz-Sarvise M, Corbo V, Tiriac H, Engle DD, Frese KK, Oni TE, et al. Identification of Resistance Pathways Specific to Malignancy Using Organoid Models of Pancreatic Cancer. Clin Cancer Res 2019;25:6742

16. Viale A, Pettazzoni P, Lyssiotis CA, Ying H, Sánchez N, Marchesini M, et al. Oncogene ablation-resistant pancreatic cancer cells depend on mitochondrial function. Nature 2014;514:628–32

17. Santana-Codina N, Roeth AA, Zhang Y, Yang A, Mashadova O, Asara JM, et al. Oncogenic KRAS supports pancreatic cancer through regulation of nucleotide synthesis. Nature Communications 2018;9:4945

18. Bryant KL, Stalnecker CA, Zeitouni D, Klomp JE, Peng S, Tikunov AP, et al. Combination of ERK and autophagy inhibition as a treatment approach for pancreatic cancer. Nat Med 2019;25:628–40

19. Kinsey CG, Camolotto SA, Boespflug AM, Guillen KP, Foth M, Truong A, et al. Protective autophagy elicited by RAF-->MEK-->ERK inhibition suggests a treatment strategy for RAS-driven cancers. Nat Med 2019;25:620–7

20. Sharma SV, Lee DY, Li B, Quinlan MP, Takahashi F, Maheswaran S, et al. A chromatin-mediated reversible drug-tolerant state in cancer cell subpopulations. Cell 2010;141:69–80

21. Wang Z, Hausmann S, Lyu R, Li TM, Lofgren SM, Flores NM, et al. SETD5-Coordinated Chromatin Reprogramming Regulates Adaptive Resistance to Targeted Pancreatic Cancer Therapy. Cancer Cell 2020;37:834–49 e13

22. Hessmann E, Johnsen SA, Siveke JT, Ellenrieder V. Epigenetic treatment of pancreatic cancer: is there a therapeutic perspective on the horizon? Gut 2017;66:168–79

23. Shen H, Laird PW. Interplay between the cancer genome and epigenome. Cell 2013;153:38–55

24. Plass C, Pfister SM, Lindroth AM, Bogatyrova O, Claus R, Lichter P. Mutations in regulators of the epigenome and their connections to global chromatin patterns in cancer. Nat Rev Genet 2013;14:765–80

25. Loewe S. The problem of synergism and antagonism of combined drugs. Arzneimittelforschung 1953;3:285–90

26. Di Veroli GY, Fornari C, Wang D, Mollard S, Bramhall JL, Richards FM, et al. Combenefit: an interactive platform for the analysis and visualization of drug combinations. Bioinformatics 2016;32:2866–8

27. Benjamini Y, Hochberg Y. Controlling the false discovery rate: a practical and powerful approach to multiple testing. J R Stat Soc Ser B Methodol 1995;57:289–300

28. Jiang H, Lei R, Ding S-W, Zhu S. Skewer: a fast and accurate adapter trimmer for next-generation sequencing paired-end reads. BMC bioinformatics 2014;15:182-

29. Patro R, Duggal G, Love MI, Irizarry RA, Kingsford C. Salmon provides fast and bias-aware quantification of transcript expression. Nat Methods 2017;14:417–9

30. Soneson C, Love MI, Robinson MD. Differential analyses for RNA-seq: transcript-level estimates improve gene-level inferences. F1000Res 2015;4:1521

31. Love MI, Huber W, Anders S. Moderated estimation of fold change and dispersion for RNA-seq data with DESeq2. Genome Biol 2014;15:550

32. Collisson EA, Sadanandam A, Olson P, Gibb WJ, Truitt M, Gu S, et al. Subtypes of pancreatic ductal adenocarcinoma and their differing responses to therapy. Nat Med 2011;17:500–3

33. Bailey P, Chang DK, Nones K, Johns AL, Patch AM, Gingras MC, et al. Genomic analyses identify molecular subtypes of pancreatic cancer. Nature 2016;531:47–52

34. Moffitt RA, Marayati R, Flate EL, Volmar KE, Loeza SG, Hoadley KA, et al. Virtual microdissection identifies distinct tumor- and stroma-specific subtypes of pancreatic ductal adenocarcinoma. Nat Genet 2015;47:1168–78

35. Subramanian A, Tamayo P, Mootha VK, Mukherjee S, Ebert BL, Gillette MA, et al. Gene set enrichment analysis: a knowledge-based approach for interpreting genome-wide expression profiles. Proc Natl Acad Sci USA 2005;102:15545–50

36. Pedersen BS, Eyring K, De S, Yang IV, Schwartz DA. Fast and accurate alignment of long bisulfite-seq reads. arXiv 2014:https://arxiv.org/abs/1401.129

37. Hansen KD, Langmead B, Irizarry RA. BSmooth: from whole genome bisulfite sequencing reads to differentially methylated regions. Genome Biol 2012;13:R83

38. Quinlan AR, Hall IM. BEDTools: a flexible suite of utilities for comparing genomic features. Bioinformatics 2010;26:841–2

39. Zerbino DR, Achuthan P, Akanni W, Amode MR, Barrell D, Bhai J, et al. Ensembl 2018. Nucleic Acids Res 2018;46:D754–D61

40. Heinz S, Benner C, Spann N, Bertolino E, Lin YC, Laslo P, et al. Simple combinations of lineage-determining transcription factors prime cis-regulatory elements required for macrophage and B cell identities. Mol Cell 2010;38:576–89

41. Roe JS, Hwang CI, Somerville TDD, Milazzo JP, Lee EJ, Da Silva B, et al. Enhancer reprogramming promotes pancreatic cancer metastasis. Cell 2017;170:875–88 e20

42. Boj SF, Hwang CI, Baker LA, Chio, II, Engle DD, Corbo V, et al. Organoid models of human and mouse ductal pancreatic cancer. Cell 2015;160:324–38

43. Li H. Aligning sequence reads, clone sequences and assembly contigs with BWA-MEM. arXiv 2013:https://arxiv.org/abs/1303.3997

44. Zhang Y, Liu T, Meyer CA, Eeckhoute J, Johnson DS, Bernstein BE, et al. Model-based analysis of ChIP-Seq (MACS). Genome Biol 2008;9:R137

45. Gaspar JM. Improved peak-calling with MACS2. bioRxiv 2018:https://doi.org/10.1101/496521

46. Leitao E, Beygo J, Zeschnigk M, Klein-Hitpass L, Bargull M, Rahmann S, et al. Locus-specific DNA methylation analysis by targeted deep bisulfite sequencing. Methods Mol Biol 2018;1767:351–66

47. Tarasov A, Vilella AJ, Cuppen E, Nijman IJ, Prins P. Sambamba: fast processing of NGS alignment formats. Bioinformatics 2015;31:2032–4

48. Garrison E, Gabor M. Haplotype-based variant detection from short-read sequencing. arXiv 2012:https://arxiv.org/abs/1207.3907

49. Köster J, Dijkstra LJ, Marschall T, Schönhuth A. Enhancing sensitivity and controlling false discovery rate in somatic indel discovery. bioRxiv 2019:https://doi.org/10.1101/741256

50. Köster J, Rahmann S. Snakemake—a scalable bioinformatics workflow engine. Bioinformatics 2012;28:2520–2

51. Li H. A statistical framework for SNP calling, mutation discovery, association mapping and population genetical parameter estimation from sequencing data. Bioinformatics 2011;27:2987–93

52. Jager M, Wang K, Bauer S, Smedley D, Krawitz P, Robinson PN. Jannovar: a java library for exome annotation. Hum Mutat 2014;35:548–55

53. R Core Team. R: A language and environment for statistical computing. Vienna, Austria: R Foundation for Statistical Computing; 2018.

54. Mazur PK, Herner A, Mello SS, Wirth M, Hausmann S, Sanchez-Rivera FJ, et al. Combined inhibition of BET family proteins and histone deacetylases as a potential epigenetics-based therapy for pancreatic ductal adenocarcinoma. Nat Med 2015;21:1163–71

55. Heid I, Steiger K, Trajkovic-Arsic M, Settles M, Eßwein MR, Erkan M, et al. Co-clinical assessment of tumor cellularity in pancreatic cancer. Clin Cancer Res 2016;23:1461–70

56. Salgia R, Kulkarni P. The Genetic/Non-genetic Duality of Drug ’Resistance’ in Cancer. Trends Cancer 2018;4:110–8

57. Notta F, Chan-Seng-Yue M, Lemire M, Li Y, Wilson GW, Connor AA, et al. A renewed model of pancreatic cancer evolution based on genomic rearrangement patterns. Nature 2016;538:378–82

58. Yin Y, Morgunova E, Jolma A, Kaasinen E, Sahu B, Khund-Sayeed S, et al. Impact of cytosine methylation on DNA binding specificities of human transcription factors. Science 2017;356

59. Xuan Lin QX, Sian S, An O, Thieffry D, Jha S, Benoukraf T. MethMotif: an integrative cell specific database of transcription factor binding motifs coupled with DNA methylation profiles. Nucleic Acids Res 2019;47:D145–D54

60. Konieczkowski DJ, Johannessen CM, Garraway LA. A convergence-based framework for cancer drug resistance. Cancer Cell 2018;33:801–15

61. Hanahan D, Coussens LM. Accessories to the crime: functions of cells recruited to the tumor microenvironment. Cancer Cell 2012;21:309–22

62. Hata AN, Niederst MJ, Archibald HL, Gomez-Caraballo M, Siddiqui FM, Mulvey HE, et al. Tumor cells can follow distinct evolutionary paths to become resistant to epidermal growth factor receptor inhibition. Nat Med 2016;22:262–9

63. McDonald OG, Li X, Saunders T, Tryggvadottir R, Mentch SJ, Warmoes MO, et al. Epigenomic reprogramming during pancreatic cancer progression links anabolic glucose metabolism to distant metastasis. Nat Genet 2017;49:367–76

64. Devarajan E, Sahin AA, Chen JS, Krishnamurthy RR, Aggarwal N, Brun A-M, et al. Down-regulation of caspase 3 in breast cancer: a possible mechanism for chemoresistance. Oncogene 2002;21:8843–51

65. Friedrich K, Wieder T, Von Haefen C, Radetzki S, Jänicke R, Schulze-Osthoff K, et al. Overexpression of caspase-3 restores sensitivity for drug-induced apoptosis in breast cancer cell lines with acquired drug resistance. Oncogene 2001;20:2749–60

66. Yang S, Zhou Q, Yang X. Caspase-3 status is a determinant of the differential responses to genistein between MDA-MB-231 and MCF-7 breast cancer cells. Biochim Biophys Acta 2007;1773:903–11

67. Bernard A, Chevrier S, Beltjens F, Dosset M, Viltard E, Lagrange A, et al. Cleaved Caspase-3 transcriptionally regulates angiogenesis-promoting chemotherapy resistance. Cancer Res 2019;79:5958–70

68. Zhou M, Liu X, Li Z, Huang Q, Li F, Li CY. Caspase-3 regulates the migration, invasion and metastasis of colon cancer cells. Int J Cancer 2018;143:921–30

69. Bestor TH, Edwards JR, Boulard M. Notes on the role of dynamic DNA methylation in mammalian development. Proc Natl Acad Sci USA 2015;112:6796

