## Supplemental_Figures for "Epigenetic plasticity via adaptive DNA hypermethylation and clonal expansion underlie resistance to oncogenic pathway inhibition in pancreatic cancer"

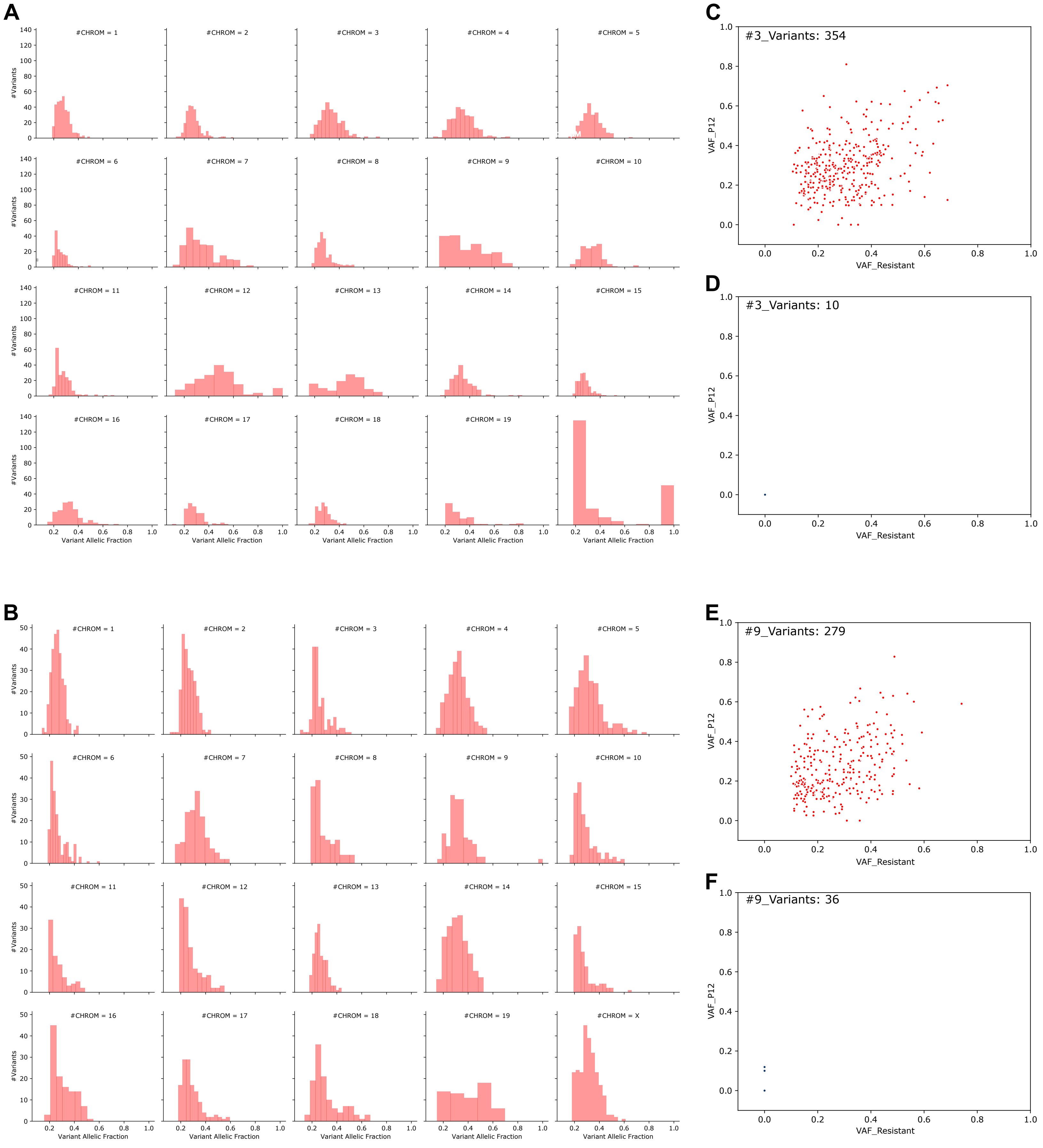


**Supplemental Fig. S1**

VAF of VpRs called by WGS separated by chromosomes for cell lines #3 **(A)** and #9 **(B)**. **(C,E)** VAF of VpRs in resistant compared to P12 cells displayed for cell lines #3 **(C)** or #9 **(E)**. **(D,F)** Scatter plot for the VAF of VpPs in resistant compared to P12 in cell lines #3 **(D)** or #9 **(F)**. Only A > T and T > A variants called by WGS and validated by WBGS are shown.

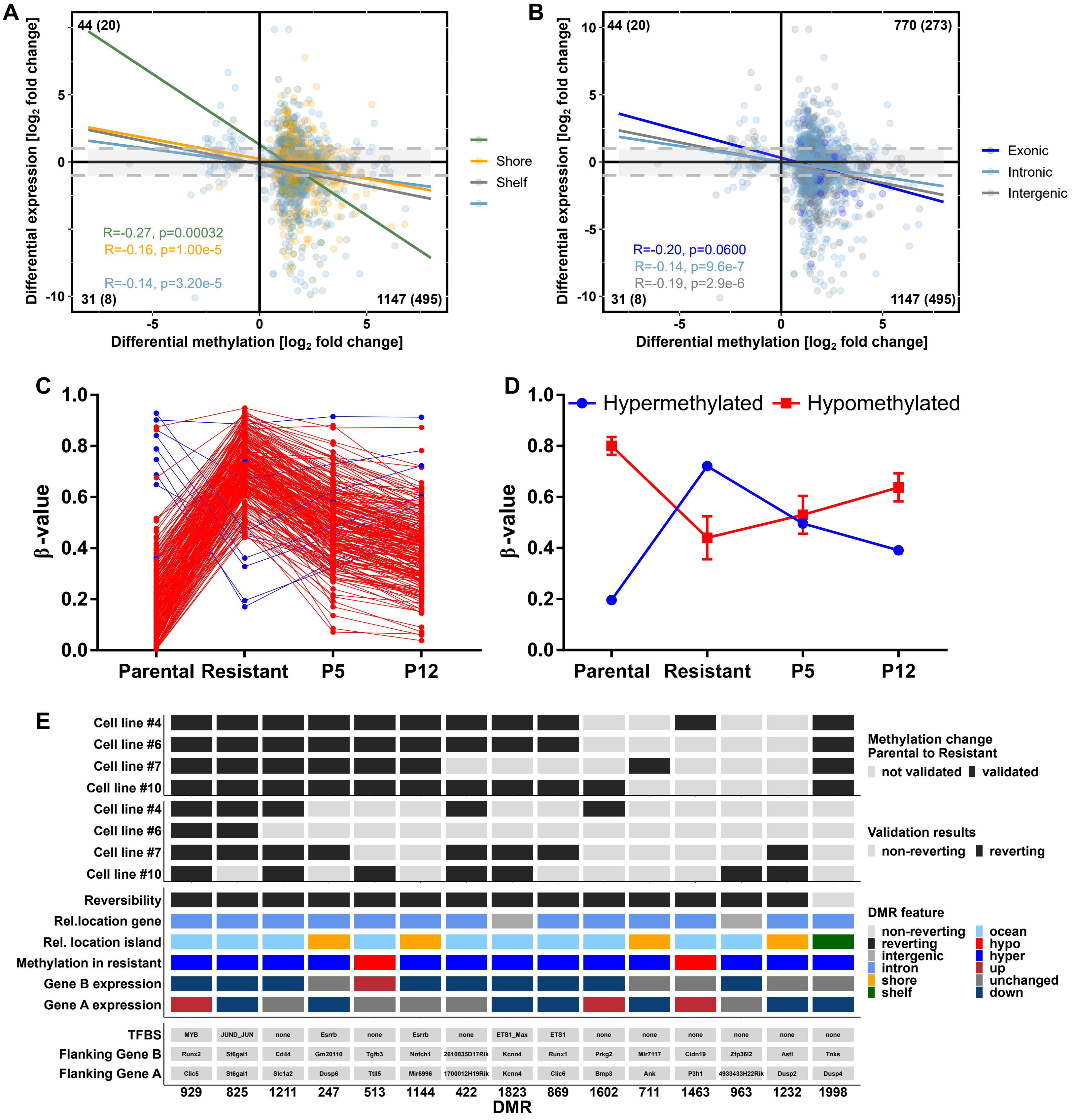


**Supplemental Fig. S2**

**(A,B)** Correlation of methylation and expression changes of DMRs between parental and resistant cells and their nearest downstream gene separated by their location according to CpG islands **(A)** or genes **(B)**. The respective Pearson correlation coefficient was calculated. Numbers indicate the number of events in the quadrant. In brackets the number of significantly differently expressed genes (adjusted p-value < 0.05 and $\log_2$ fold change < -1 or > 1) are displayed. Dotted grey lines mark $\log_2$ fold changes of 1 or -1. **(C)** Methylation pattern of 217 reverting DMRs divided into hypo- (blue) and hypermethylated (red) regions. **(D)** Mean DNA methylation of 217 reverting DMRs ± SEM separated into hypo- and hypermethylated regions. **(E)** Validation results of 15 DMRs analyzed by targeted deep bisulfite sequencing in four independent cell lines. In addition, features and annotations are displayed for each DMR.

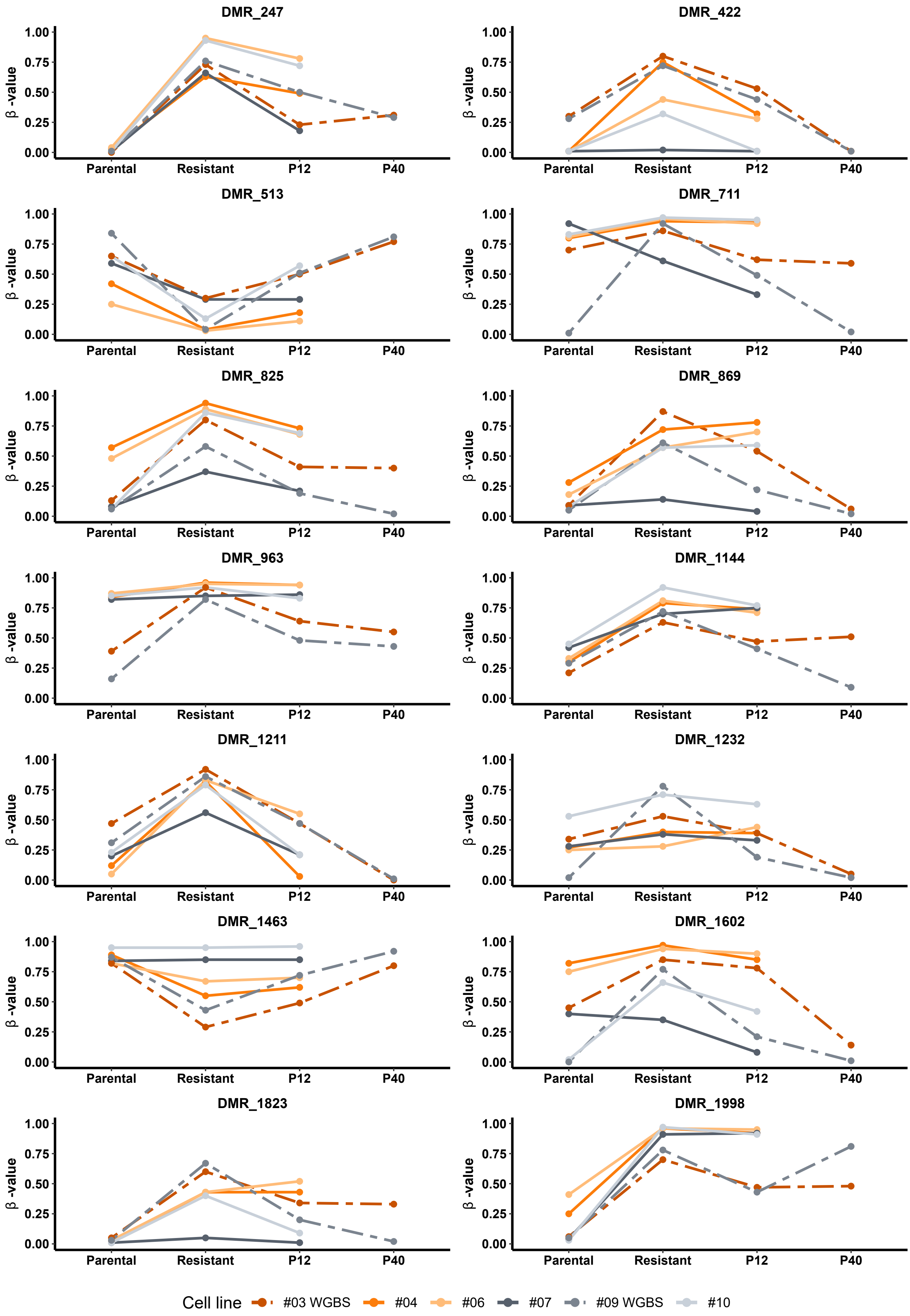


**Supplemental Fig. S3**

Individual methylation patterns of 14 DMRs validated by targeted deep bisulfite sequencing in four independent cell lines compared to cell lines #3 and #9 analyzed by WGBS. In addition, P40 of #3 and #9 was measured by targeted deep bisulfite sequencing. DMR_929 is displayed in Fig. 4D,E.

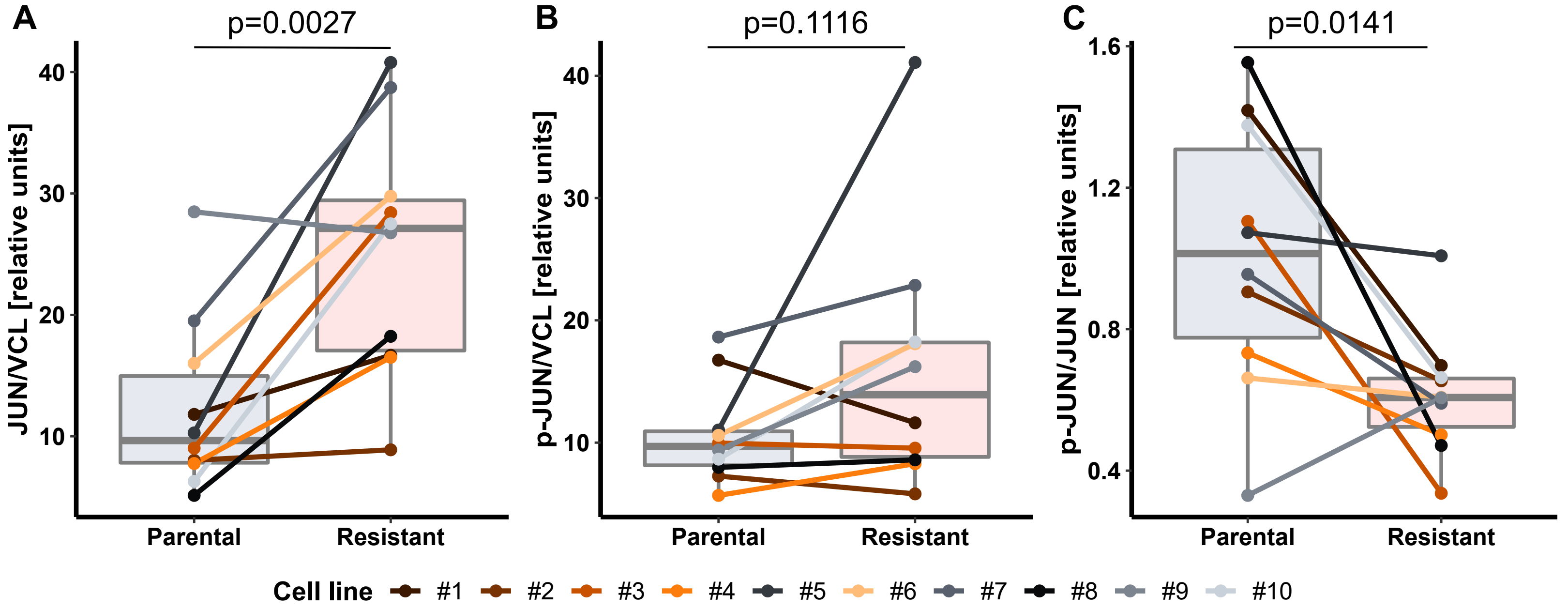


**Supplemental Fig. S4**

Relative protein expression of total JUN **(A)** or p-JUN **(B)** compared to VCL in parental and resistant cells. **(C)** Proportion of p-JUN to JUN. Statistics were calculated using the two-tailed paired Student's t-test.

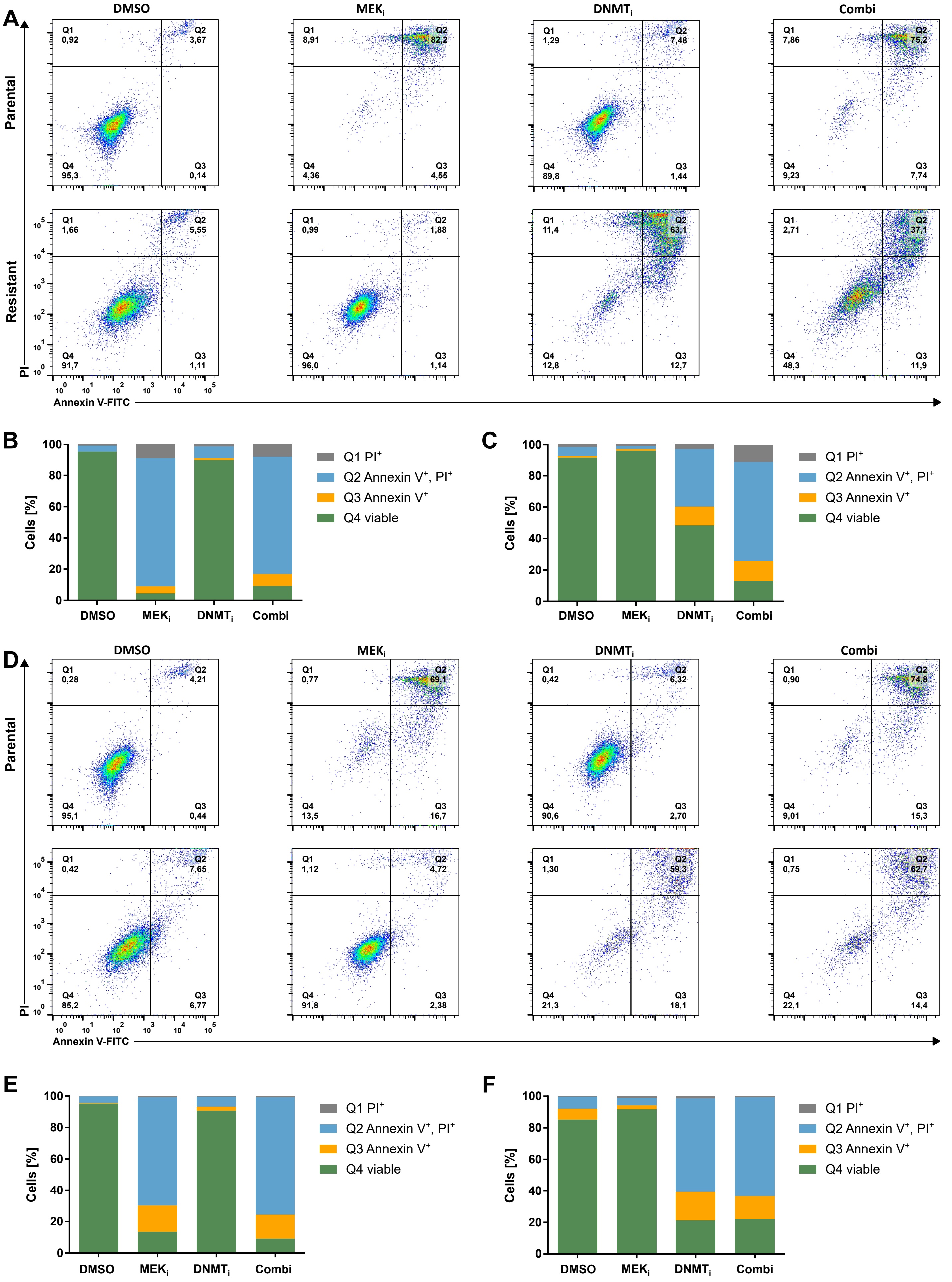


**Supplemental Fig. S5**

Annexin V-FITC/PI co-staining measured by FACS after 84 h of indicated treatment. **(A-C)** Dot plots **(A)** and quantification for cell line #3 parental **(B)** and resistant **(C)**. **(D-F)** Dot plots **(D)** and quantification for cell line #4 parental **(E)** and resistant **(F)**.

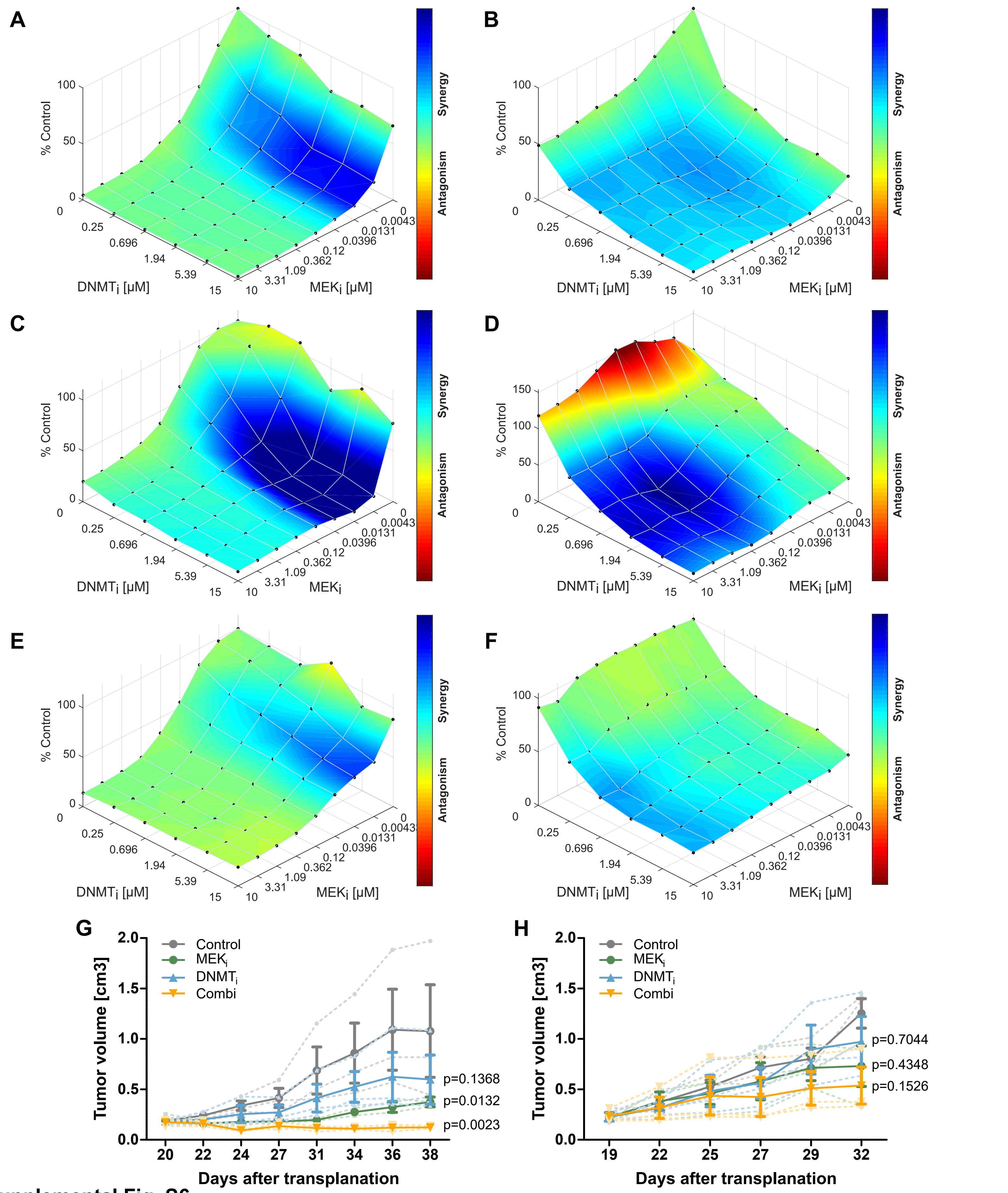


### Supplemental Fig. S6

**(A-F)** Synergy analysis of MEK$_i$ plus DNMT$_i$ using the Loewe method of the Combenefit software shown for cell lines #4 **(A,B)**, #9 **(C,D)** and #10 **(E,F)**. Results for parental cells are shown left, while resistant cells are displayed in the right panels, respectively. **(G,H)** Two different PDX of PDAC treated either with MEK$_i$, DNMT$_i$ or the combination. Solid lines represent the mean tumor volume of the three mice per treatment group ± SEM.
