## Supplemental_Tables_S1_S3_S6 for "Epigenetic plasticity via adaptive DNA hypermethylation and clonal expansion underlie resistance to oncogenic pathway inhibition in pancreatic cancer"

**Supplemental Table S1**

Primer sequences used for targeted deep bisulfite sequencing of selected differentially methylated regions (DMRs). During oligonucleotide synthesis the following tag sequences were added to the 5´end of each forward and reverse primer: CTTGCTTCCTGGCACGAG (forward) and CAGGAAACAGCTATGAC (reverse)

| DMR | Primer | Sequence | PCR product location in mm10 |
|---|---|---|---|
| 247 | forward | ATTTTATGGGTTAGTTGTTTGATTT | chr10:99266158-99266419 |
| | reverse | CAACTAATAACACTTTTAACACCC | |
| 422 | forward | GTTTGTTTGTAATTGGGGATTAGG | chr11:113060998-113061290 |
| | reverse | TCTAAAATACAACCCTACATATAAATC | |
| 513 | forward | TTGTTAAATGGAATTGTTGTTGAGA | chr12:85859330-85859634 |
| | reverse | AAAACTCTTACATAAACTACCAAAC | |
| 711 | forward | TAAAGTAGTGGGATAAATTTTTTTT | chr15:27467984-27468252 |
| | reverse | CTAATCTTCTTATTCTTAACAAATCC | |
| 825 | forward | AGATTAGTGTTTTTAATTATTTTGA | chr16:23272009-23272213 |
| | reverse | CCAAACAAAAAACATCATACTTTC | |
| 869 | forward | TTTTTTGTTTAAGGAAAGGATA | chr16:92620511-92620772 |
| | reverse | CCATTCTCAAAACAAATTTTAC | |
| 929 | forward | TTGTAAGGTTGTATTTTTTTTGATGTTT | chr17:44718034-44718321 |
| | reverse | CAACCCTTAAATACTAAAACTCAACTC | |
| 963 | forward | TGGGAGGTAGTGTGGGATATAGTAG | chr17:84141583-84141880 |
| | reverse | CTTCCCAAAAACAAAAACACTCTAA | |
| 1144 | forward | AGGAGTGTGTTAATTTTTAGGGGTTA | chr2:26501667-26501948 |
| | reverse | CCCAACCAAATAAAACCTACCTAA | |
| 1211 | forward | ATTTGTTAGTATAGAAGAAGGTTGGTAGT | chr2:102873250-102873543 |
| | reverse | AACAAATTCTAAAAAAAATTCCCTC | |
| 1232 | forward | TTTGTTTAGGTTTTGTTTTTTGTTG | chr2:127336828-127337044 |
| | reverse | ATTCACACCATTTACAAATCACAC | |
| 1463 | forward | GTTGATTATTAGTTTTTTTTGTAGTT | chr4:119244934-119245233 |
| | reverse | CACATTTACACTAATATCCCAACC | |
| 1602 | forward | TAGTAATATAAGATGTTTAAGTATTGAA | chr5:98943301-98943589 |
| | reverse | AAAACCCCACAAAACTCTCTCTAA | |
| 1823 | forward | AAAAGTTTTAGAGTGAGTAGAATAGGT | chr7:24370317-24370596 |
| | reverse | CTTACAACCCAAAAACCACAAC | |
| 1998 | forward | TAATTTTAGTATATGGGGAGTGGGA | chr8:34813921-34814236 |
| | reverse | TAACCATTCCCTTAAAAACACTACC | |

**Supplemental Table S2**

IC50 of trametinib in the parental cells based on averaged dose response curves.

| Cell line | IC50 [nM] | Replicates |
|---|---|---|
| #1 | 25.2 | 2 |
| #2 | 9.4 | 2 |
| #3 | 10.9 | 3 |
| #4 | 11.2 | 3 |
| #5 | 12.9 | 2 |
| #6 | 16.3 | 3 |
| #7 | 14.3 | 3 |
| #8 | 5.3 | 2 |
| #9 | 55.2 | 3 |
| #10 | 10.9 | 3 |

**Supplemental Table S6**

Numbers (No.) of evaluated mitosis are displayed (NA, not available).

| No._mitosis | Karyotype_cell line #3 | Karyotype_cell line #9 |
|---|---|---|
| 1 | 76 | 70 |
| 2 | 79 | 70 |
| 3 | 78 | 70 |
| 4 | 74 | 67 |
| 5 | NA | 59 |
| 6 | NA | 70 |
| 7 | NA | 70 |
| 8 | NA | 60 |
| 9 | NA | 73 |
| 10 | NA | 70 |
| 11 | NA | 63 |
| 12 | NA | 70 |
